# Solar UVB Radiation for the Synthesis of Vitamin D_3_ in Four Skin Types – Model Propagation Studies

**DOI:** 10.1101/2024.12.04.626916

**Authors:** Pradeep K. Pasricha, Arun K. Upadhayaya

## Abstract

The study presents model computations of the wavelength of absorption of radiation energy in the ultraviolet (UV) band in the epidermis cells keratinocytes for four skin types: White-skin, Pale Yellow-skin, Brown-skin and Black-skin. Keratinocytes cells are idealized as spherical dielectric microcavities, with central melanin granule and a layer of keratin molecules surrounding the cell. Refractive indices of epidermis tissue, melanin and keratin are obtained from the published data. The size distribution of keratinocytes in the epidermis is obtained with the conjecture that the differentiated keratinocytes grow exponentially in the process of cell differentiation. The mole concept is used to calculate the size of melanin granules, one each for four skin types, using the published data of melanin content of the four skin types. The thickness of keratin layer around the cell keratinocyte does not occur in the model computations. In White-skin, the model solar UVB radiation wavelength range of 296 – 300 nm is close to the maximal range of 295 – 300 nm, of the range 290 – ∼317 nm, of the synthesis of vitamin D3 by the compound 7-dehydrocholesterol in skin cells. In Pale Yellow-skin, Brown-skin and Black-skin, the model solar UVB/(limited UVA) radiation wavelength ranges for the synthesis of vitamin D_3_ are 290 – 295 nm, 290 – 297 nm and 307 – 314 nm, and 313 – ∼317 nm, respectively. The solar UVB radiation synthesizes vitamin D_3_ to the maximum in White-skin and the least in Black-skin. The synthesis of vitamin D_3_ is to a lesser extent in Pale Yellow-skin compared to White-skin. The solar UVB radiation synthesizes ample levels of vitamin D_3_ in the pigmented Brown-skin. The solar UVB radiation is more effective to synthesize vitamin D_3_ in Pale Yellow-skin compared to Brown-skin, since the Brown-skin is prone to pigmentation. The UVA radiation wavelength ranges that cause tanning of skin are analyzed. In Brown-skin, in particular, the wavelength range is 315 – 342 nm. Also, the physical mechanism of the UVA-caused tanning is similar to that of the UVB-caused tanning of skin. The UVA radiation phototherapies toward tanning of skin may be applied in the wavelength range of 340 – 370 nm. A sunscreen gel, in Brown-skin, in particular, may have the formulation to absorb the effective solar UVB and UVA radiation of wavelengths of 290 – 310 nm and 315 – 340 nm, respectively. The UVB radiation phototherapies to synthesize vitamin D_3_, especially for individuals allergic to medicinal vitamin D_3_, have varied model wavelength ranges for the four skin types. The study emphasizes the specific role of UV radiation toward synthesis of vitamin D_3_ and therapeutic use in skin tanning for different skin types. The preparation of a sunscreen gel for skin protection may vary with skin type.

## 1. Introduction

The Sun’s visible exterior region of photosphere at a temperature of about 6000 K emits optical radiation, that is, electromagnetic waves in a broad spectrum of ultraviolet, visible and infrared wavelengths. The electromagnetic spectrum of ultraviolet (UV) radiation in the wavelength range 100 – 400 nm is categorized into three bands, UVC (100 – 280 nm), UVB (280 – 315 nm) and UVA (315 – 400 nm). The visible radiation wavelength band is: 400 – 700 nm. The near-infrared radiation wavelength band is: 700 – 1400 nm. The infrared radiation wavelengths are above 700 nm up to 1 mm (= 10^6^ nm). The solar radiation UV, visible and near-infrared primarily penetrates the human skin. The solar radiation shorter than about 290 nm, pertaining to UVC (and lower wavelengths) and part of UVB, fail to reach the earth’s surface since these are absorbed by the atmosphere. The main atmospheric constituents are nitrogen (N_2_), oxygen (O_2_), water vapor (H_2_O) and carbon dioxide (CO_2_). (Water vapor and carbon dioxide are minor constituents, that is, of tiny amounts.) The overall atmospheric constituents absorb the solar radiation as quanta (photons) of energy hν (ν = c/λ, ν the frequency of radiation, λ the wavelength of radiation and h the Planck’s constant). The radiation energy (hν) is taken up in photo-ionization, photo-dissociation, photo-chemical reactions and formation of minor constituent ozone (O_3_). Thus there are produced the ionized region ionosphere and the ozone region in the atmosphere. The ozone so formed absorbs the UVC radiation and the UVB radiation to 97%, and more, of its level at the top of the atmosphere. The absorption spectrum of ozone depicts a steady decline in absorption of UVB radiation to negligible extent in the wavelength range 280 to 315 nm [1, Figure 27 B]. The increased absorption is due to the higher path length through the ozone region, as the amount of absorbed photons increase with the consequent higher absorbed energy (J, joule). The solar zenith angle, measured with respect to the vertical axis, determines the angle of the sun at the latitude of the place on the earth. It is minimal when the sun is nearly overhead at the local noon. The amount of UVB radiation reaching the earth’s surface depends on latitude, altitude, time, season, and etcetera. (The solar radiation is intense during the summer, with longer brighter days, though the earth’s orbit is ‘around aphelion’, that is, farther away from the sun. The solar radiation is less intense during the winter though the earth’s orbit is ‘around perihelion’, that is, nearer to the sun.) The UVA radiation reaches the earth’s surface almost unaffected by the atmosphere. The places within the tropics (latitudes within about 23.5º N and 23.5º S around the equator) receive intense UV radiation since rays of sun fall upon it straight, and not inclined. (As a consequence, the inhabitants of tropics are of dark skin.) There are a number of environmental parameters that determine the extent of solar UV radiation reaching the earth’s surface; thin cloud cover allows transmission of solar UV radiation [2, 3]. The visible radiation is reduced to almost 50% on the earth’s surface. The visible radiation is absorbed by the green pigment chlorophyll in plants due to photochemical reactions, that is, the process of photosynthesis [4, page 1091]. (Plants, green algae, etcetera, make oxygen for human respiration during photosynthesis. Ocean is the other source of oxygen.) The photosynthesis takes place with the absorption of solar visible radiation energy (hν) that enables water (H_2_O) to combine with carbon dioxide (CO_2_) to form carbohydrates and oxygen. The organic compounds carbohydrates are eventually broken down into its components in plants, releasing energy for the plants to grow and gain mass. Some of the released energy is converted to heat (enthalpy, ΔH). (Plants during the process of respiration release carbon dioxide.) The infrared (heat) radiation on the earth’s surface is around 40% of its level at the top of the atmosphere. Infrared radiation results in the sensation of warmth on the skin. Near-infrared radiation finds applications in medical sciences, in particular dermatology [5]. The excessive exposure to solar UV radiation leads to sun-tanning (darkening) and sunburn (reddening, erythema) of the skin, and skin cancer. The use of sunscreen gels protects the skin from exposure to UV radiation. A sunscreen gel with Sun Protection Factor (SPF)-50 absorbs approximately 95% of solar UVB radiation. (Sunscreen gels contain organic (aromatic) compounds that absorb UV radiation energy and convert it to heat (IR radiation) [4, page 650].) The use of sunscreen gels proportionally reduces the synthesis of vitamin D_3_ in the skin. The UV therapy (phototherapy) includes exposure of the UVB radiation on the skin to enhance vitamin D_3_ levels in humans. The UVA exposure on the skin (therapeutic use in skin care) is used for the tanning of the skin.

The top skin layer epidermis (tissue, thickness ∼ 0.115 mm, ∼ 115 µm; 1 µm = 10^−3^ mm) mainly comprises the basal layer (the inner layer, the stratum basale, thickness ∼ 0.01 mm, ∼ 10 µm) and the stratum corneum (the outer layer, the skin surface, the keratin layer, thickness ∼ 0.015 mm, ∼ 15 µm). The middle layer dermis of the skin (thickness ∼ 1 mm, ∼ 1000 µm) lies beneath the epidermis. The thickness of various layers cited is as per the skin model adopted by Zhang et al. [6]. (Therein the epidermis without the stratum corneum is classified as the living epidermis (thickness ∼ 0.1 mm). The stratum corneum has a thickness of ∼ 0.015 mm. The living epidermis overlies the dermis (thickness ∼ 1 mm).) The layer stratum spinosum, above the basal layer, is close to the stratum corneum. (There are two more layers in between the stratum spinosum and the stratum corneum, namely, stratum granulosum and stratum lucidum.) The basal layer is adjacent to the dermis. The elastic connective tissue in the dermis holds capillary blood vessels, nerves, and etcetera. The bottom layer hypodermis of the skin lies beneath the dermis. The subcutaneous tissue in the hypodermis mainly consists of cells that store fat.

The basal layer of the epidermis contains the stem cells and the cells melanocytes. The stem cells produce the cells keratinocytes. The cells melanocytes generate the organelles melanosomes within the cells. The pigment of skin, an organic colouring compound, melanin, is synthesized and contained in melanosomes. The melanosomes are subsequently transferred to nearby keratinocytes, and the melanin is poured out in the keratinocyte’s cytoplasm [7, Review]. In the basal layer a keratinocyte divides to give a daughter cell, with an increase in the cell number, that is, the cell proliferation. A daughter cell is not subject to division, and the cells do not further proliferate. (It is akin to an atom, that is, nucleus of atom, which disintegrates in a radioactive substance; the daughter nucleus is not subject to disintegration.) The cell proliferation represents the “birth” of the cells keratinocytes. About 90% of the cells in the epidermis (the skin cells) are keratinocytes. The size of the proliferated keratinocytes cells is in the range 5 to 11 µm, being predominant in the range 5 to 7 µm [8, 9]. (The thickness of the basal layer may be taken as the size of the proliferated keratinocytes ∼ 10 µm, ∼ 0.01 mm.) Beyond the basal layer, the proliferated keratinocytes cells in the epidermis enter into the process of cell differentiation, where the keratinocyte cells steadily evolve in size up to skin surface. These differentiated keratinocytes in the epidermis undergo terminal differentiation, that is, the decay (death) of the cells, at the skin surface. In the process the pigment melanin reaches the skin surface that determines the skin pigmentation, the skin color. (Keratinocytes in the epidermis are said to undergo terminal differentiation, that is, they migrate (“move outward”) toward the skin surface carrying pigment melanin with them and decay.) The cytoplasm of the decayed (dead) keratinocytes is keratin. The decayed (dead) keratinocytes, the corneocytes, in the keratin layer (the surface layer, the stratum corneum) are shed off, and are replenished from within the basal layer in about 28 days. The size of a corneocyte is approximately 30 to 50 µm in diameter and 1 µm thick [10]. Thus the “birth-to-decay” span of the keratinocytes is every 28 days; the proliferated keratinocytes in the basal layer are reproduced spontaneously every day. The jelly-like cellular material in the epidermis becomes thicker, harder and flattened keratin in the stratum corneum, which protects the body from the outside environment.

The solar UVB radiation is absorbed in the epidermis layer. The longer solar UVA wavelengths penetrate the dermis layer. The exposure to solar UVB and UVA radiation leads to tanning of the skin, the dark skin. The UVA radiation initiates the tanning of the skin immediately upon exposure to the sun. The normal exposure to the UVA radiation leads to tanning of the skin for a few hours. The above normal exposure to the UVA radiation leads to tanning of the skin for a day, or more number of days, and diminishes to the ambient normal (of genetic origin) skin pigmentation. The “immediate” short-term UVA-caused tanning of the skin is due to the chemical modifications of the organic compound melanin. The UVA range of wavelengths that leads to tanning of the skin has a peak between 340 nm to 370 nm [7, Review]. The skin tanned by the UVA exposure does not protect the skin from the UVB-caused sunburn (erythema). The tanning of the skin by the excessive exposure to the UVB radiation appears after a delay of a number of days of the exposure, and lasts for two weeks to three weeks [7, Review]. The UVB-caused “delayed” long-term tanning of the skin is due to the greater production of pigment melanin (melanin mass density, ng cm^-3^) by the melanocytes, with the appearance of enhanced skin pigmentation. The UVA range of wavelengths 320 nm to 360 nm is also effective, though to a lesser extent, in stimulating the production of pigment melanin and the skin pigmentation [3]. The tanning caused by the UVB/UVA(320 – 360 nm) protects the skin from further harmful over exposure to the UV radiation, that is, sunburn (erythema). The pigment melanin absorbs the solar UVB radiation, which is needed to synthesize vitamin D_3_ in the epidermis layer of the skin. (The solar UVB radiation of wavelengths 280 – 290 nm does not reach the earth’s surface. The absorption spectrum of pigment melanin is 290 – 700 nm, with higher absorption at the UV wavelengths. The UV radiation reaches the epidermis (up to the dermis) about four times less in dark skin compared to light skin [7, Review].) The sunlight is the natural source of vitamin D_3_. The organic compound Vitamin D occurs as vitamin D_2_ (calciferol) and vitamin D_3_ (cholecalciferol). Vitamin D_2_ is obtained from diet. Vitamin D_3_ is also obtained in the medicinal form as cholecalciferol capsules and sachets of cholecalciferol granules. The synthesis of vitamin D_3_ is partly due to photochemical reactions, the photosynthesis, with the absorption of solar UVB radiation energy (hν) by the compound 7-dehydrocholesterol (provitamin D_3_) in the skin cells. (The photosynthesis by 7-dehydrocholesterol is in part akin to the photosynthesis by the green pigment chlorophyll of plants [4, page 1091]. The photosynthesis by the green pigment in plants occurs in cell organelles chloroplasts. Heat (enthalpy, ΔH) is released during photosynthesis.) The compound 7-dehydrocholesterol is found mostly in the epidermis layer of the skin, with the highest concentrations in the basal layer and the stratum spinosum [11]. Thus the synthesis of vitamin D_3_ occurs in these layers of the epidermis, which almost occupy the entire epidermis. The compound 7-dehydrocholesterol in the skin, the cutaneous (referring to the skin) 7-dehydrocholesterol, absorbs the UVB – UVA radiation of wavelengths 280 nm to ∼317 nm to form previtamin D_3_, with the release of heat (enthalpy, ΔH) [1, Figure 10]. From Figure 10 of Reference 1 of Wacker and Holick, the epidermal absorption spectrum of the UVB – UVA radiation in the wavelengths 280 – ∼317 nm reveals a roughly bell-shaped curve with maximal absorption in the wavelengths 295 – 300 nm, peaked around 298 nm. The absorption by the cutaneous 7-dehydrocholesterol is minimal around 317 nm. Also, the absorption by the cutaneous 7-dehydrocholesterol around 280 nm is similar to that around 305 nm. The effective absorption of solar UVB by the cutaneous 7-dehydrocholesterol may be taken to be in the wavelengths 290 – 310 nm. Previtamin D_3_ in the skin is transformed to vitamin D_3_ using the released heat ΔH, a process known as thermal isomerization. Since the solar UVB radiation of wavelengths 280 – 290 nm does not reach the earth’s surface, the epidermis layer of the skin absorbs the solar UVB – UVA radiation of wavelengths 290 – ∼317 nm to synthesize vitamin D_3_ in the skin. Vitamin D_3_ in the skin is transported to the body through the capillary blood vessels in the dermis [1]. (The sunlight synthesized-previtamin D_3_ in the skin is converted to vitamin D_3_ to one hundred percent, compared to orally ingested-vitamin D_3_ of sixty percent [1].)

In the present study, model computations of the propagation of electromagnetic waves in the epidermis of four types of skin are performed (Section 6). The four skin types are: (a) White-skin, (b) Pale Yellow-skin, (c) Brown-skin and (d) Black-skin. The propagation wavelengths are the UVB and UVA ultraviolet bands. For the model computations, an idealized spherical differentiated keratinocyte of a given diameter (say, 15 µm) in the epidermis of (say, White-skin) is considered. The pigment melanin consists of granules of uniform shape for the given skin type. A spherical melanin granule of a given diameter (say, 0.8 µm, White-skin) is at the center of the idealized keratinocyte. The spherical keratinocyte is surrounded by a layer of keratin. Thus a spherical dielectric microcavity is considered, which consists of a core of melanin and a pair of layers of keratinocyte and keratin. The thickness of keratin layer does not occur in the computations. Electromagnetic waves of given wavelength λ (nm, say, in UVB band) are incident in the core melanin. The wavelength of resonance of the dielectric microcavity enables to determine the wavelength of propagation in the idealized keratinocyte in the epidermis. The calculations in the UVB band are performed to one-decimal-place in wavelength (δλ = 0.1 nm), where a range of wavelengths of 3 nm is considered as the wavelengths of propagation. The calculations in the UVA band are performed to one-place in wavelength (δλ = 1 nm), where a range of wavelengths of 5 nm is considered as the wavelengths of propagation. The model computations are carried out for a number of diameters of keratinocytes in the epidermis of a given skin type with a single melanin granule diameter. The size distribution of differentiated keratinocytes in the epidermis is obtained with the conjecture that they evolve exponentially with time in the epidermis (Section 4). The differentiated keratinocytes evolve from size ∼ 5 to 7 µm just above the basal layer to size ∼ 30 to 50 µm in the stratum corneum, where they are eliminated as corneocytes in 28 days. The granules of pigment melanin are considered to be molecules of uniform shape in different skin types. The size (nm) of melanin granules is calculated using the published data of melanin content (mmol dl^-1^, millimole per deciliter) for the four skin types (Section 2). Refractive indices of epidermis tissue, melanin and keratin (in the stratum corneum) appropriate to UVB and UVA bands are used from the published data (Section 5). The model computed (model) wavelengths of propagation in the UVB band are analyzed in reference to (a) solar radiation and (b) phototherapies toward synthesis of vitamin D_3_ in the epidermis of different skin types. And, in the UVA band, these are analyzed in reference to (c) solar radiation and (d) phototherapies (therapeutic use in skin care) toward tanning of skin. Also, in the UVB and UVA bands, the model computed (model) wavelengths are analyzed for (e) preparation of sunscreen gel for different skin types.

## 2. Size of Melanin Granule and Melanin Density From Melanin Content

The melanin content is obtained by making measurements of reflectance spectra on the back of ten volunteers using diffuse reflectance spectroscopy [12]. The published data for four types of skin having ten values of melanin content (× 10^−7^ mmol per dl, mmol dl^-1^, millimole per deciliter; 1 dl = 100 cm^3^) are the following: (a) Caucasian – 2.7, 4.5, (b) Japanese – 14, 15, 17, 24, (c) Brown – 35, 47, 67 and (d) black African – 118. The average values of melanin content are (a) 3.5 × 10^−7^ mmol dl^-1^, (b) 17 × 10^−7^ mmol dl^-1^, (c) 50 × 10^−7^ mmol dl^-1^ and (d) 118 × 10^−7^ mmol dl^-1^, respectively. (In Reference 12, Specific Brown-skin is not mentioned; let it be Indian.) The four skin types are herewith referred to as (a) White-skin, (b) Pale Yellow-skin, (c) Brown-skin and (d) Black-skin, respectively. The forms of pigment melanin of pheomelanin and eumelanin are found in White-skin and Black-skin, respectively.

The pigment melanin is a polymeric organic compound formed from repeating subunits monomers. There are 28 monomer units and 46 monomer units in pheomelanin and eumelanin, respectively [13]. The average monomer molar masses of pheomelanin and eumelanin are 274 g mol^-1^ (gram per mole) and 171 g mol^-1^ (gram per mole), respectively. Thus, the molar masses of pheomelanin and eumelanin are 7672 (274 x 28 = 7672) g mol^-1^ and 7866 (171 x 46 = 7866) g mol^-1^, respectively. The melanin polymer may be represented as a granule of melanin (pheomelanin, eumelanin) of molecular mass (weight) M_w_ (Da, dalton, g mol^-1^, gram per mole) and size d (nm; nanometer, 1 nm = 10^−9^ m).

In the following calculations, the melanin granule (polymer) is considered to be a molecule of uniform shape, whose size d (nm) may be estimated from the melanin content.

Let the melanin content be X moles cm^-3^.

1 cm^3^ of melanin has X moles. 1 mole contains 6.02 × 10^23^ melanin polymeric molecules (granules). (The Avogadro constant, N_A_ = 6.02 × 10^23^ mol^-1^, is the number of atoms or molecules per mole, by the mole concept.)

The number of granules N that occupy a volume of 1 cm^3^ is

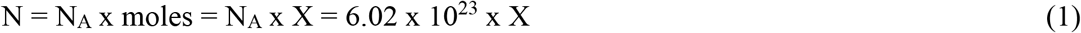

The volume per granule V (cm^3^) is

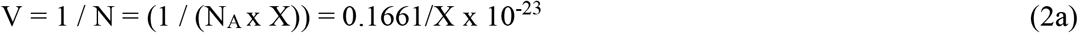

The volume per granule V (nm^3^; 1 cm = 10^7^ nm; 1 cm^3^ = 10^21^ nm^3^) is

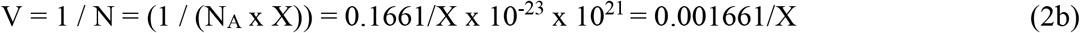

The volume occupied per granule is of the order of d^3^, where d (nm) is the size of granule (polymer).

From (2b),

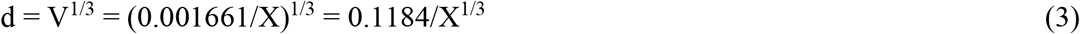

The mass density ρ (g cm^-3^) of melanin (pheomelanin, eumelanin) of molecular mass M_w_ (Da) is calculated. The melanin content is X moles cm^-3^.

The molecular mass of M_w_ (Da, g mol^-1^) implies that 1 mole of melanin has a mass of M_w_ g. So, the mass density ρ (g cm^-3^) of melanin is given by

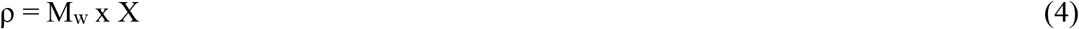

The number density of melanin granules N (cm^-3^) is given by (1).

The expression of particle size d (nm) of a substance of molecular mass M_w_ (Da) and density ρ (g cm^-3^) is given as [14]

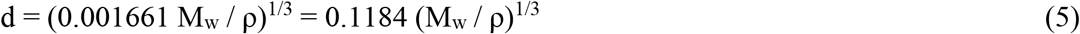

where 0.001661 is (1/6.02) × 10^−2^. 6.02 is part of the Avogadro number. From (5), the mass density ρ (g cm^-3^) is

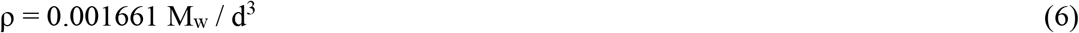

### 2.1 Size of Melanin Granule and Melanin Density in White-Skin

The measured melanin (pheomelanin) content in White-skin is X = 3.5 × 10^−7^ mmol dl^-1^ (1dl = 100 cm^3^). The molecular mass of pheomelanin is M_w_ = 7672 g mol^-1^.

1 dl has 3.5 × 10^−7^ millimole of melanin.

So, 100 cm^3^ has 3.5 × 10^−7^ × 10^−3^ mole of melanin.

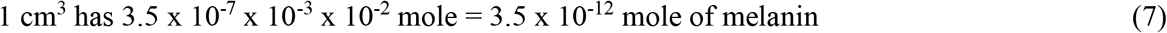

The granule size d (nm), from (3), is

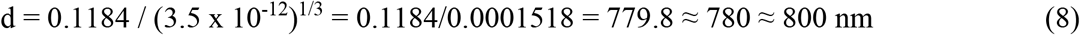

The mass density of melanin ρ (g cm^-3^), from (4) and (7), is

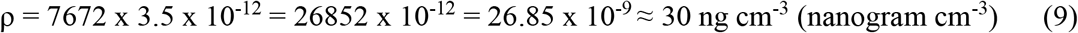

The granule number density N (cm^-3^), from (1) and (7), is

N = 6.02 × 10^23^ x 3.5 × 10^−12^ ≈ 2 × 10^12^ cm^-3^ = 0.2 × 10^13^ cm^-3^

One may verify the value of the granule size d (nm), from (5) and (9), with M_w_ = 7672 g mol^-1^, as

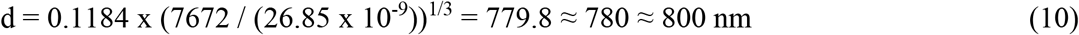

The values of d (nm) in (8) and (10) are in tally.

### 2.2 Size of Melanin Granule and Melanin Density in Black-Skin

The measured melanin (eumelanin) content in Black-skin is X = 118 × 10^−7^ mmol dl^-1^. The molecular mass of eumelanin is M_w_ = 7866 g mol^-1^. 1 dl (= 100 cm^3^) has 118 × 10^−7^ millimole of melanin. 1 cm^3^ has 118 × 10^−12^ mole of melanin.

The granule size d (nm) is d = 0.1184 / (118 × 10^−12^)^1/3^ = 241.4 ≈ 240 nm

The mass density of melanin ρ (g cm^-3^) is ρ = 7866 x 118 × 10^−12^ = 928.2 × 10^−9^ ≈ 930 ng cm^-3^ (nanogram cm^-3^)

The granule number density N (cm^-3^) is N = 6.02 × 10^23^ x 118 × 10^−12^ ≈ 70 × 10^12^ cm^-3^ = 7 × 10^13^ cm^-3^

### 2.3 Size of Melanin Granule and Melanin Density in Brown-Skin

The measured melanin content in Brown-skin is X = 50 × 10^−7^ mmol dl^-1^. 1 dl (= 100 cm^3^) has 50 × 10^−7^ millimole of melanin. 1 cm^3^ has 50 × 10^−12^ mole of melanin.

The granule size d (nm) is d = 0.1184 / (50 × 10^−12^)^1/3^ = 321.4 ≈ 320 nm

The granule number density N (cm^-3^) is N = 6.02 × 10^23^ x 50 × 10^−12^ ≈ 30 × 10^12^ cm^-3^ = 3 × 10^13^ cm^-3^

### 2.4 Size of Melanin Granule and Melanin Density in Pale Yellow-Skin

The measured melanin content in Pale Yellow-skin is X = 17 × 10^−7^ mmol dl^-1^. 1 dl (= 100 cm^3^) has 17 × 10^−7^ millimole of melanin. 1 cm^3^ has 17 × 10^−12^ mole of melanin.

The granule size d (nm) is d = 0.1184 / (17 × 10^−12^)^1/3^ = 460.5 ≈ 460 nm

The granule number density N (cm^-3^) is N = 6.02 × 10^23^ x 17 × 10^−12^ ≈ 10 × 10^12^ cm^-3^ = 10^13^ cm^-3^

### 2.5 A Summary and Size of Melanin Granule for Model Studies of Wave Propagation in Epidermis

A summary of melanin content X (mmol dl^-1^), granule size d (nm), melanin mass density ρ (ng cm^-3^) and granule number density N (cm^-3^) for the four skin types is given in Table 1. The White-skin and Black-skin have pheomelanin and eumelanin, respectively. The molecular mass M_w_ (Da) in Pale Yellow-skin and Brown-skin are unknown. Hence, the melanin mass density in these skin types is not calculated.

**Table 1.** Summary of melanin content (mmol dl^-1^), granule size (nm), melanin mass density (ng cm^-3^) and granule number density (cm^-3^) for the four skin types, viz., White-skin, Pale Yellow-skin, Brown-skin and Black-skin. The melanin content is obtained from [12]

| Skin Type | Melanin Content<br>X (mmol dl<sup>-1</sup>) | Melanin Granule<br>Size<br>d (nm) | Melanin Mass<br>Density<br>$\rho$ (ng cm<sup>-3</sup>) | Granule Number<br>Density<br>N (cm<sup>-3</sup>) |
| --- | --- | --- | --- | --- |
| White-skin | $3.5 \times 10^{-7}$ | 800 | 30 | $0.2 \times 10^{13}$ |
| Pale Yellow-skin | $17 \times 10^{-7}$ | 460 | – | $10^{13}$ |
| Brown-skin | $50 \times 10^{-7}$ | 320 | – | $3 \times 10^{13}$ |
| Black-skin | $118 \times 10^{-7}$ | 240 | 930 | $7 \times 10^{13}$ |

The melanin granule size for White-skin, Pale Yellow-skin, Brown-skin and Black-skin are 800 nm, 460 nm, 320 nm and 240 nm, respectively. These are used for the model studies of propagation of UVB and UVA waves in the epidermis of the four skin types (Section 6). The refractive index of melanin in the UVB and UVA bands is tabulated in Section 5.

## 3. Number Density of Keratin Molecules in Stratum Corneum and Its Comparison with Number Density of Skin Pigmentation

The keratin layer (stratum corneum) is made up of dead keratinocytes whose cytoplasm is keratin. Keratin is an organic compound with a molecular mass (M_w_) of about 50 kDa [15, 16] and mass density (ρ) of 1.3 g cm^-3^ [17]. The size (d, nm) of the keratin molecule in the keratin layer, considering it to be a molecule of uniform shape, may be calculated from (5), d = 0.1184 x (50 × 10^3^ / 1.3)^1/3^ = 0.1184 x 33.75 ≈ 4 nm

One may calculate the number density of keratin molecules as follows. The molecular mass of keratin M_w_ = 50000 Da implies, 1 mole of keratin has a mass of 5 × 10^4^ g = 50000 g.

1 cm^3^ of keratin has ρ/M_w_ = 1.3/50000 = 0.26 × 10^−4^ moles.

1 mole contains the Avogadro number N_A_ = 6.02 × 10^23^ of keratin molecules (by the mole concept).

So, N = N_A_ x moles = N_A_ x ρ/M_w_ = 6.02 × 10^23^ x 0.26 × 10^−4^ ≈ 1.5 × 10^19^ molecules occupy a volume of 1 cm^3^. Thus, the number density of keratin molecules, that is, per unit volume of stratum corneum, is 1.5 × 10^19^ cm^-3^.

One may estimate the number of pigment granules (size ∼ 400 nm) corresponding to the number of keratin molecules (size ∼ 4 nm), per unit volume of stratum corneum, as: 1.5 × 10^19^ x 4^3^ / 400^3^ ≈ 10^13^ melanin granules. The estimated number density of pigment granules matches with the skin pigmentation (granules) of size ∼ 400 nm, having a number density ≈ 10^13^ cm^-3^, in Table 1 of Section 2.

A layer of keratin is used in the model studies of propagation of UVB and UVA waves in the epidermis of the aforementioned four skin types (Section 6). The refractive index of keratin in the UVB and UVA bands is tabulated in Section 5. The thickness of the keratin layer is not required in the model propagation computations.

## 4. Size Distribution of Keratinocytes in Epidermis

In the phenomenon of cell differentiation in the epidermis, the keratinocytes evolve in size from the basal layer to the stratum corneum. The main factor that promotes cell differentiation is the steady increase in the concentration of alkaline-earth element calcium (Ca) in the epidermis. The concentration of calcium increases from the basal layer to the stratum corneum, with the highest concentration in the stratum granulosum [18]. (Stratum granulosum is the nearest layer to the stratum corneum.) The concentration of calcium is high in the stratum corneum. Vitamin D_3_ prompts calcium uptake in the body. The supposed to be a linear system the cell differentiation may be described by the mathematical model using the first-order differential equation in two steps. First, the keratinocytes in the epidermis undergo terminal differentiation (that is, the decay) at the skin surface. They are shed off from the skin surface every 28 days. It enables to calculate the decay of keratinocytes per day, the decay constant (day^-1^). Secondly, the decay constant (day^-1^) may be used to calculate the progressive increase in the size of the keratinocytes in the epidermis.

In the cell differentiation in the epidermis, the number of differentiated keratinocytes that in a given time enter into differentiation may be described by the first-order differential equation (11). The proliferated keratinocytes of size 5 µm to 11 µm in the basal layer are reproduced spontaneously every day; the predominant proliferated size range is 5 µm to 7 µm. Upon terminal differentiation (that is, the decay) every 28 days, the size of a differentiated keratinocyte is approximately 30 to 50 µm. In order to calculate the decay constant (day^-1^), it is adopted that not all the keratinocytes decay, but un-decayed keratinocytes amounting to 1%, 5% or 10% are the remnants at the skin surface every 28 days.

The first-order differential equation that describes the decay of keratinocytes is given by

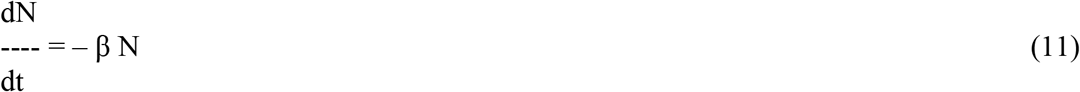

where N is the number of keratinocytes present in the epidermis (without the stratum corneum) at time t and β the decay constant of the keratinocytes. (The decay constant is conventionally denoted by the Greek alphabet lambda (λ). However, it is denoted by the Greek alphabet beta (β) since the symbol λ represents the wavelength of waves in the present study. In a radioactive substance, the first-order equation describes the number of atoms (that is, nuclei of atoms) that disintegrate per second. The decay constant β is the probability that the atom will decay in one second.) The solution of differential equation (11) is given by

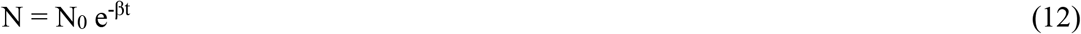

where N_0_ is the number of keratinocytes present in the epidermis (at the basal layer) at time t = 0. The number N of keratinocytes in the epidermis decreases exponentially with time. From (12),

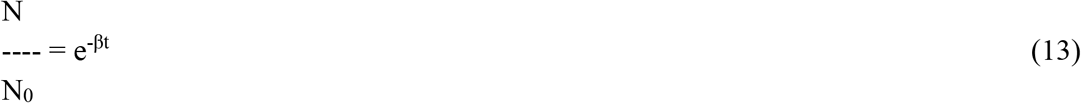

The decay constant β equals the fraction of keratinocytes that decay per unit time (say, per day, day^-1^).

The decay constant β (day^-1^) for the decay of keratinocytes, for the three cases of un-decayed keratinocytes of (i) N/N_0_ = 1%, (ii) N/N_0_ = 5% and (iii) N/N_0_ = 10% at the skin surface every 28 days, are obtained.

i. Decay constant β for 1% un-decayed keratinocytes at the skin surface In (13), N/N_0_ is 1% (= 0.01). It implies, by taking natural logarithms of both sides, – 4.6 = – β t With the decay period of keratinocytes of t = 28 days, β = 0.164 day^-1^
ii. Decay constant β for 5% un-decayed keratinocytes at the skin surface In (13), N/N_0_ is 5% (= 0.05). It implies, by taking natural logarithms of both sides, – 3.0 = – β t With the decay period of keratinocytes of t = 28 days, β = 0.107 day^-1^
iii. Decay constant β for 10% un-decayed keratinocytes at the skin surface In (13), N/N_0_ is 10% (= 0.1). It implies, by taking natural logarithms of both sides, – 2.3 = – β t With the decay period of keratinocytes of t = 28 days, β = 0.082 day^-1^

An estimation of the size distribution of keratinocytes in the epidermis (without the stratum corneum) may be obtained with the conjecture that the size of the differentiated keratinocytes A (µm) grows exponentially with time, with constant (–)β (day^-1^), in the process of cell differentiation. In view of the first-order differential equation (11), the size of differentiated keratinocytes decreases exponentially from the skin surface, as do the number of keratinocytes beyond the basal layer, with the decay constant β (day^-1^). Thus, one has

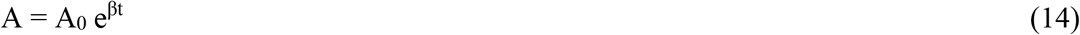

where A_0_ (µm) is the size of proliferated (undifferentiated) keratinocytes in the basal layer. From (14), the size distribution of differentiated keratinocytes in the epidermis is obtained, with the proliferated keratinocytes of size 5 µm, 7 µm, 9 µm and 11 µm and decay constant 0.164 day^-1^, 0.107 day^-1^ and 0.082 day^-1^, seven days apart over 28 days.

### 4.1 Size Distribution of Keratinocytes with Un-Decayed Keratinocytes of 1% at the Skin Surface

For the un-decayed keratinocytes of 1% at the skin surface over 28 days, the decay constant β equals 0.164 day^-1^. The keratinocytes almost entirely decay at the skin surface over 28 days. Using (14), Table 2 depicts the size A (µm) of differentiated keratinocytes in the epidermis for the size A_0_ (µm) of proliferated keratinocytes of 5 µm to 11 µm, seven days apart over 28 days. The differentiated keratinocytes are of large sizes over 21 days. The size of keratinocytes, underlined in Table 2, is more than the thickness of the epidermis (∼ 0.1 mm, ∼ 100 µm) for the range of sizes of proliferated keratinocytes of 5 µm to 11 µm.

**Table 2.** Size (µm) of differentiated keratinocytes 7 days apart over 28 days for the un-decayed keratinocytes at the skin surface of 1%. The size of proliferated keratinocytes is 5 µm to 11 µm 5. The underlined keratinocyte sizes are more than the thickness of the epidermis (∼ 0.1 mm, ∼ 100 µm)

| Time apart<br>(day) | Differentiated<br>Keratinocyte size<br>A (μm)<br>(Proliferated<br>Keratinocyte size<br>A<sub>0</sub> = 5 μm) | Differentiated<br>Keratinocyte size<br>A (μm)<br>(Proliferated<br>Keratinocyte size<br>A<sub>0</sub> = 7 μm) | Differentiated<br>Keratinocyte size<br>A (μm)<br>(Proliferated<br>Keratinocyte size<br>A<sub>0</sub> = 9 μm) | Differentiated<br>Keratinocyte size<br>A (μm)<br>(Proliferated<br>Keratinocyte size<br>A<sub>0</sub> = 11 μm) |
| --- | --- | --- | --- | --- |
| 0 | 5 | 7 | 9 | 11 |
| 7 | 16 | 22 | 28 | 35 |
| 14 | 50 | 70 | 89 | $\approx 100$ |
| 21 | <u>157</u> | <u>219</u> | <u>282</u> | <u>344</u> |
| 28 | <u>493</u> | <u>690</u> | <u>888</u> | <u>1086</u> |

### 4.2 Size Distribution of Keratinocytes with Un-Decayed Keratinocytes of 5% at the Skin Surface

For the un-decayed keratinocytes of 5% at the skin surface over 28 days, the decay constant β equals 0.107 day^-1^. The differentiated keratinocytes decay to 95% at the skin surface over 28 days. Using (14), Table 3 depicts the size A (µm) of differentiated keratinocytes in the epidermis for the size A_0_ (µm) of proliferated keratinocytes of 5 µm to 11 µm, seven days apart over 28 days. The differentiated keratinocytes are of sizes between 5 µm to 100 µm over 28 days for the proliferated keratinocyte size of 5 µm. The differentiated keratinocytes are of large sizes over 28 days for the size of proliferated keratinocytes of 7 µm to 11 µm. The size of keratinocytes, underlined in Table 3, is more than the thickness of the epidermis (∼ 0.1 mm, ∼ 100 µm).

**Table 3.**
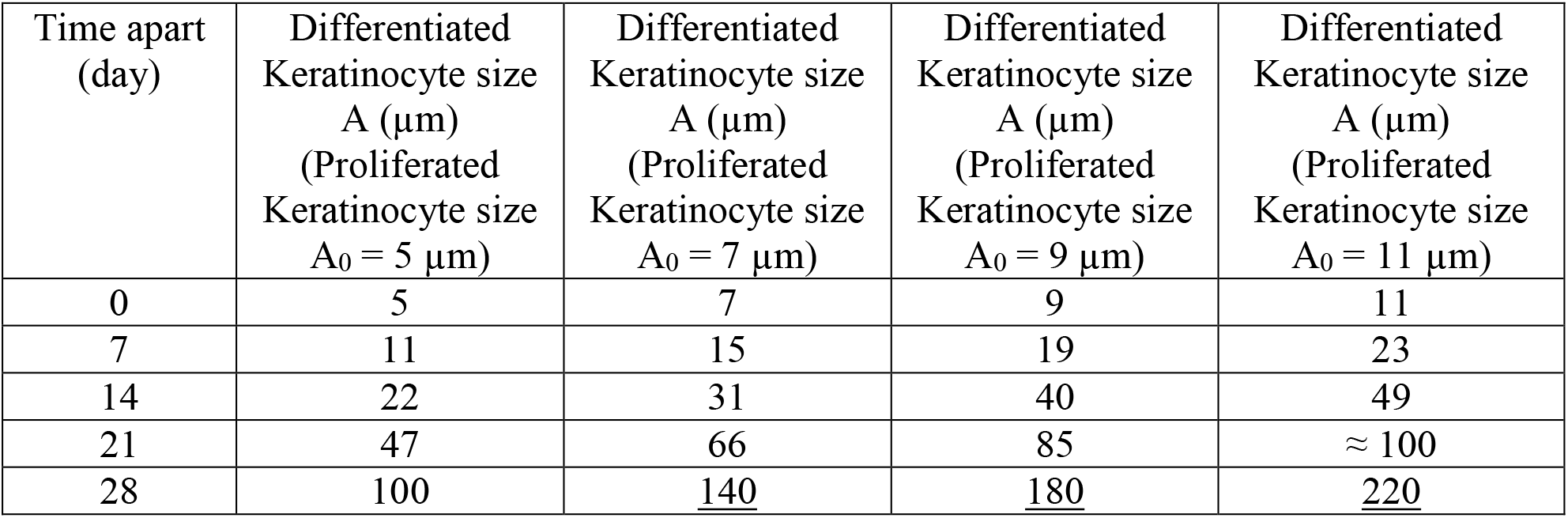
Size (µm) of differentiated keratinocytes 7 days apart over 28 days for the un-decayed keratinocytes at the skin surface of 5%. The size of proliferated keratinocytes is 5 µm to 11 µm. The underlined keratinocyte sizes are more than the thickness of the epidermis (∼ 0.1 mm, ∼ 100 µm)

| Time apart<br>(day) | Differentiated<br>Keratinocyte size<br>A ($\mu\text{m}$)<br>(Proliferated<br>Keratinocyte size<br>A<sub>0</sub> = 5 $\mu\text{m}$) | Differentiated<br>Keratinocyte size<br>A ($\mu\text{m}$)<br>(Proliferated<br>Keratinocyte size<br>A<sub>0</sub> = 7 $\mu\text{m}$) | Differentiated<br>Keratinocyte size<br>A ($\mu\text{m}$)<br>(Proliferated<br>Keratinocyte size<br>A<sub>0</sub> = 9 $\mu\text{m}$) | Differentiated<br>Keratinocyte size<br>A ($\mu\text{m}$)<br>(Proliferated<br>Keratinocyte size<br>A<sub>0</sub> = 11 $\mu\text{m}$) |
| --- | --- | --- | --- | --- |
| 0 | 5 | 7 | 9 | 11 |
| 7 | 11 | 15 | 19 | 23 |
| 14 | 22 | 31 | 40 | 49 |
| 21 | 47 | 66 | 85 | $\approx 100$ |
| 28 | 100 | <u>140</u> | <u>180</u> | <u>220</u> |

**Table 4.**
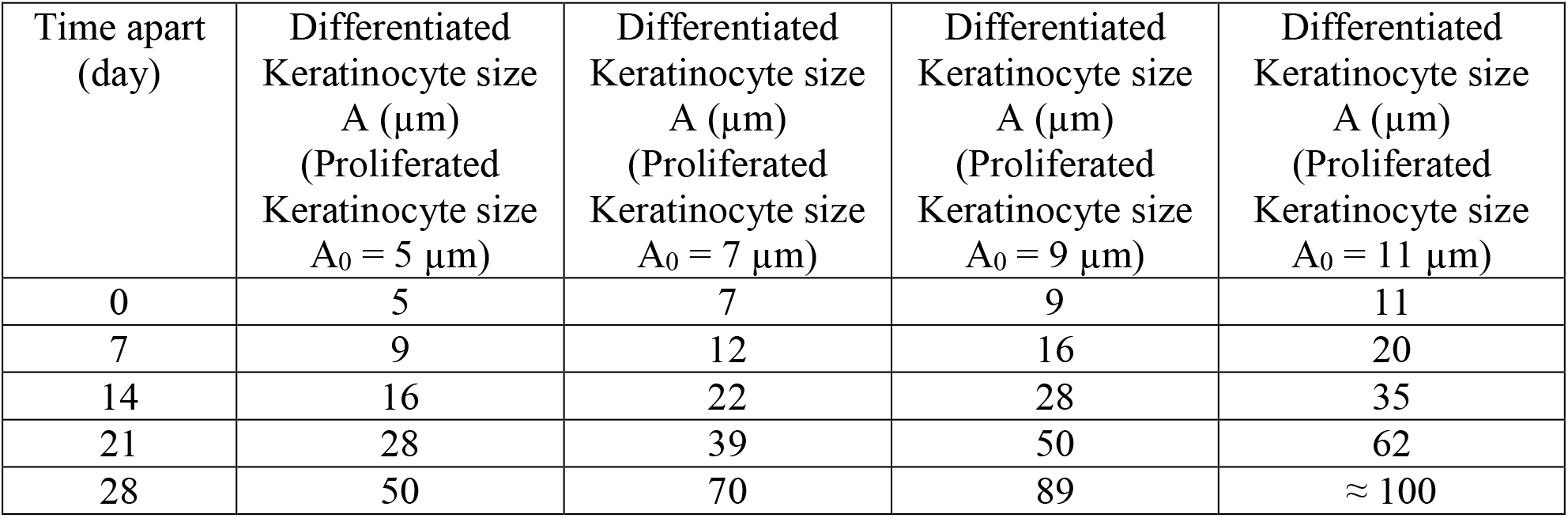
Size (µm) of differentiated keratinocytes 7 days apart over 28 days for the un-decayed keratinocytes at the skin surface of 10%. The size of proliferated keratinocytes is 5 µm to 11 µm

For the un-decayed keratinocytes of 7% at the skin surface over 28 days, the decay constant β equals 0.095 day^-1^. The differentiated keratinocytes decay to 93% at the skin surface over 28 days. The differentiated keratinocytes are of sizes between 5 µm to 70 µm over 28 days for the proliferated keratinocyte size of 5 µm. Also, the differentiated keratinocytes are of sizes between 7 µm to 100 µm over 28 days for the proliferated keratinocyte size of 7 µm. The size of differentiated keratinocytes is between about 15 µm to 50 µm for both the proliferated keratinocytes of size 5 µm and 7 µm, over day-7 – to – day-21.

### 4.3 Size Distribution of Keratinocytes with Un-Decayed Keratinocytes of 10% at the Skin Surface

For the un-decayed keratinocytes of 10% at the skin surface over 28 days, the decay constant β equals 0.082 day^-1^. The differentiated keratinocytes decay to 90% at the skin surface over 28 days. Using (14), Table 3 depicts the size A (µm) of differentiated keratinocytes in the epidermis for the size A_0_ (µm) of proliferated keratinocytes of 5 µm to 11 µm, seven days apart over 28 days. The differentiated keratinocytes are of sizes between 5 µm to 70 µm over 28 days for the proliferated keratinocyte sizes of 5 µm and 7µm. The differentiated keratinocytes are of sizes between 5 µm to 100 µm over 28 days for the range of sizes of proliferated keratinocytes of 5 µm to 11µm. The size of differentiated keratinocytes is between about 15 µm to 50 µm over 28 days for the proliferated keratinocytes of size 5 µm to 11 µm, over day-7 – to – day-21.

### 4.4 A Summary and Size of Keratinocytes for Model Studies of Wave Propagation in Epidermis

To summarize: (i) The size of differentiated keratinocytes lies between about 10 µm to 70 µm for the proliferated keratinocytes of size 5 µm to 7 µm, over day-0 – to – day-28. (ii) The size of differentiated keratinocytes lies between about 15 µm to 70 µm for the proliferated keratinocytes of size 5 µm to 11 µm, over day-7 – to – day-21.

The differentiated keratinocyte size of 15 µm, 30 µm, 50 µm and 70 µm are used for the model studies of propagation of UVB and UVA waves in the epidermis of the aforementioned four skin types (Section 6). The keratinocyte of diameters of 15 µm to 70 µm correspond to the epidermis as the keratinocytes evolve in size toward the skin surface. The refractive index of epidermis tissue in the UVB and UVA bands is tabulated in Section 5.

## 5. Refractive Indices of Epidermis Tissue, Melanin and Keratin in the UVB and UVA bands

The refractive indices of epidermis tissue, melanin and keratin in stratum corneum, appropriate to various wavelength bands, are obtained from the published data.

The complex refractive index of the epidermis tissue, for the Caucasian (White-skin) and African American (Black-skin) types, is measured at eight wavelengths between 325 nm and 1557 nm [19, Figure 6]. Let the complex refractive index be n = (n_r_ + in_i_). The real part n_r_ is the index of refraction. The imaginary part n_i_ signifies absorption (or scattering) of the optical radiation in the medium. In the epidermis tissue, it is the scattering of the optical radiation by the turbid medium (tissue) that results in the reduction of the intensity of optical radiation. The refractive index of the epidermis tissue (n = n_r_), in column (1) of Table 5, is obtained from the smoothened curve of n_r_ versus wavelength λ (nm) between 325 and 800 nm for the two types of skin [19, Figure 6]. The isolated “peaks” in the imaginary refractive index (n_i_) in wavelength bands, in Table 5, are significantly correlated with high melanin mass density (ng cm^-3^) in Black-skin type [19]. A smoothened curve of n_i_ versus wavelength λ (nm) is also obtained between 325 and 800 nm [19, Figure 6]. The imaginary refractive index n_i_ in column (2) of Table 5 depicts these indices for the White-skin and Black-skin types. The authors of Reference 19, Ding et al., conclude that the high melanin mass density (ng cm^-3^) in Black-skin significantly affects the imaginary refractive index (for instance, n_i_ ≈ 0.0150 at λ = 442 nm, Table 6). As an extension of the present study, the smoothened values of refractive index (n_r_) versus wavelength (λ, nm) are fitted with the Cauchy dispersion relation by the method of non-linear least-squares (Appendix A). (As an additional illustration, the refractive index of gold nanoparticles of size 10 to 15 nm at a wavelength of 500 nm is (1 + i2), with n_r_ = 1 and n_i_ = 2. The intensity of an incident beam of optical radiation is reduced due to absorption in such a medium.)

**Table 5.** Refractive indices (n) of epidermis tissue, melanin and keratin appropriate to UVB, UVA, visible and near-infrared bands. n_r_ is the real part of refractive index of epidermis tissue. “Peaks” in the imaginary part of refractive index (n_i_) of epidermis tissue and “Maxima” in refractive index (n) of melanin are at depicted at the corresponding wavelengths for the given skin types

| Wavelength Bands ($\lambda$) | Refractive Index ($n = n_r$) of Epidermis Tissue [19]<br>(1) | Peaks in Refractive Index ($n_i$) of Epidermis Tissue [19]<br>(2) | Refractive Index ($n$) of Melanin [20]<br>(3) | Maxima in Refractive Index ($n$) of Melanin [20]<br>(4) | Refractive Index ($n$) of Keratin [21, 6]<br>(5) |
| --- | --- | --- | --- | --- | --- |
| UVB<br>(280 – 315 nm) | 1.49<br>($\lambda = 325$ nm) | – | 1.69<br>($\lambda = 300$ nm) | 1.70 – 1.73 – 1.70<br>($\lambda = 285 – 300 – 315$ nm) | 1.55 |
| UVA<br>(315 – 400 nm) | 1.48 | 0.0075<br>($\lambda = 325$ nm)<br>(White-skin) | 1.67 | 1.68<br>($\lambda = 350 – 400$ nm) | 1.55 |
| Visible<br>(400 – 475 nm)<br>(475 – 700 nm) | 1.45<br>1.43 | 0.0150<br>($\lambda = 442$ nm)<br>(White-skin, Black-skin)<br>0.0100<br>($\lambda = 532$ nm)<br>(White-skin, Black-skin)<br>0.0125<br>($\lambda = 633$ nm)<br>(White-skin) | 1.65<br>1.63 | 1.68<br>($\lambda = 400 – 450$ nm)<br>1.67<br>($\lambda = 525 – 575$ nm) | 1.55<br>1.55 |
| Near-infrared<br>(700 – 800 nm) | 1.43 | 0.0125<br>($\lambda = 850$ nm<br>$\approx 800$ nm)<br>(White-skin) | 1.62 | | 1.54 |

**Table 6.** The wavelengths of resonance of spherical dielectric microcavity representing melanin embedded keratinocyte of size 15 µm to 70 µm in the epidermis of White-skin in the UVB and UVA bands. The condition of resonance is termed as “good” (“g”) or “fair” (“f”). A range of 3 nm in the UVB and 5 nm in the UVA is considered as the wavelengths of resonance

| Wavelength Bands ($\lambda$) | White-skin Keratinocyte Size 15 $\mu\text{m}$ | White-skin Keratinocyte Size 30 $\mu\text{m}$ | White-skin Keratinocyte Size 50 $\mu\text{m}$ | White-skin Keratinocyte Size 70 $\mu\text{m}$ |
| --- | --- | --- | --- | --- |
| UVB (280 – 315 nm) | 296.4 nm (“f”)<br>–<br>296.5–299.8 nm (“g”)<br>–<br>299.9 nm (“f”) | 280.0–283.6 nm (“f”) | 284.2–287.1 nm (“g”) | 280.0–283.0 nm |
| UVA (315 – 400 nm) | 361–371 nm (“g”) | 317–320 nm (“f”)<br>–<br>321–326 nm (“g”)<br>355–362 nm (“g”) | 353–356 nm (“g”)<br>–<br>357–359 nm (“f”) | 352–355 nm (“g”)<br>–<br>356–357 nm (“f”) |

The organic compound polyindoledione is a form of melanin. Polyindoledione and melanin have almost identical optical properties [20]. The refractive index of polyindoledione, and thus of melanin, is measured between 200 nm to 800 nm. The refractive index of melanin (n), in column (3) of Table 5, is obtained from the smoothened curve of n versus wavelength λ (nm) between 250 and 800 nm in Figure 2 of Repenko et al. [20]. The broad “maxima” in the refractive index (n) in wavelength bands are depicted in column (4) of Table 5. For the visible range of wavelengths in Table 5, the wavelengths of “peaks” in the imaginary part of refractive index n_i_ for the Black-skin tend to match the wavelengths of “maxima” in the refractive index of melanin: λ = 442 nm (column (2)) and λ = 400 – 450 nm (column (4)) and λ = 532 nm (column (2)) and λ = 525 – 575 nm (column (4)). The presence of melanin is also evident in the visible range 460 – 560 nm of the diffuse spectra, observed between 460 – 800 nm, from four types of skin [12]. The visible range of wavelengths 400 – 500 nm is also effective in tanning of the skin [7, Review]. As an extension of the present study, the smoothened values of refractive index (n) versus wavelength (λ, nm) are fitted with the Cauchy dispersion relation by the method of non-linear least-squares (Appendix A).

The refractive index of keratin in the stratum corneum, in column (5) of Table 5, is n = 1.55 at λ = 400 nm to 700 nm [21, 6]. The refractive index of keratin, in column (5) of Table 5, is n = 1.54 at λ = 700 nm to 800 nm [6].

The refractive indices of epidermis tissue, melanin and keratin, appropriate to various wavelength bands, are given in column (1), column (3) and column (5), respectively, of Table 5. These are used for the model studies of propagation of UVB and UVA waves in the epidermis of the aforementioned four skin types (Section 6). Table 5 provides the refractive indices in the spectrum of UVB, UVA, visible (400 – 700 nm) and part of near-infrared (700 – 800 nm) bands. The refractive indices in the visible and near-infrared bands are intended for further model studies of wave propagation in the epidermis of human skin.

## 6. Model Studies of Propagation of UVB and UVA Waves in the Epidermis of White-Skin, Pale Yellow-Skin, Brown-Skin and Black-Skin

Model computations of the propagation of electromagnetic waves in a differentiated keratinocyte in the epidermis of human skin are performed. The wavelength bands are the UVB (280 – 315 nm) and UVA (315 – 400 nm). For the model computations, an idealized spherical keratinocyte in the epidermis is considered. The idealized keratinocyte has four distinct diameters (sizes), that is, (i) 15 µm (15000 nm), (ii) 30 µm (30000 nm), (iii) 50 µm (50000 nm) and (iv) 70 µm (70000 nm) (Section 4). The keratinocyte of diameter of 15000 nm corresponds to the epidermis in the vicinity of the basal layer. The keratinocyte of diameters of 30000 nm to 70000 nm correspond to the epidermis, away from the basal layer, as the keratinocytes evolve in size (“migrate”) toward the skin surface. The idealized keratinocyte has at its center a spherical melanin granule of a given diameter (size). Four diameters of melanin granules are considered: 800 nm (White-skin), 460 nm (Pale Yellow-skin), 320 nm (Brown-skin) and 240 nm (Black-skin) (Section 2). A layer of keratin surrounds the keratinocyte. The layer thickness of keratin around the keratinocyte does not occur in the model computations. Thus, there is formed a spherical microcavity, which consists of a core of melanin and a pair of layers of keratinocyte and keratin. The refractive indices (n) of the dielectric medium of the microcavity, that is, melanin, epidermis tissue and keratin, are given in Table 5 of Section 5. The electromagnetic waves of wavelength λ (nm) are incident in the core melanin. The wavelength of resonance of the dielectric microcavity at the wavelength λ (nm) is examined. Thereby, the wavelength λ (nm) of the propagation of electromagnetic waves is computed.

A spherical microcavity is termed as a Bragg Reflector [22], and also a Bragg Onion Resonator without a stem [23]. (A stem is an opening, which is used to excite TE/TM modes of monochromatic electromagnetic waves. The formulation of a Bragg Onion Resonator of Xu et al. [23] for TE_10_ (and TM_10_ mode) of wave propagation has been coded in the C–programming language [24a, 24b, 24c]. In the present computations, the C programming code has been used to compute the wavelength of resonance of the spherical microcavity representing the idealized keratinocyte. (A similar set of computations can be made for TM_10_ mode.) One has to compute the “final” complex matrix T (2 x 2) at a number of wavelengths λ_i_ (nm), i = 1, 2… 100 (say), at an increment of δλ = 1 nm (say). Then, for the occurrence of resonance of the microcavity, one has to examine the condition on the complex matrix elements T_21_ ≈ –T_22_ at each wavelength λ_i_ (nm).

In the present study, the condition of resonance at a given wavelength λ (nm) is termed as: “high” (“h”) or “good” (“g”) or “fair” (“f”). For illustration, in a microcavity consisting of core of ZnTe (n = 2.7) and eight pairs of layers of ZnTe (n = 2.7) and SiO_2_ (n = 1.45) (Section 6.1), the occurrence of resonance at λ = 1385 nm is “high” (“h”), with the following elements of the complex matrix T (2 x 2):

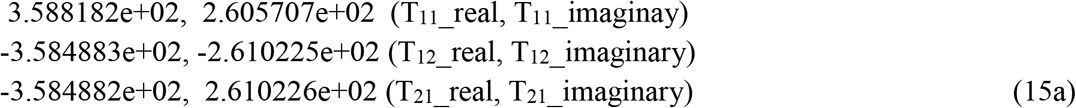

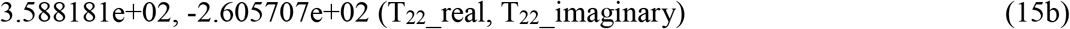

It is the actual output of the aforementioned computer program. Here, “e” stands for the base-10. For instance, e+02 is the number 10^02^. Let the complex numbers in (15a) and (15b) be given by T_21_ ≡ (–a + ib) and T_22_ ≡ (c – id), respectively, in the rectangular system of coordinates. The condition of occurrence of “high” (“h”) resonance implies that a ≈ c and b ≈ d. In terms of matrix elements, the condition of occurrence of resonance is: T_21__real ≈ – T_22__real (i.e., –3.584882e+02 ≈ (–) 3.588181e+02) and T_21__imaginary ≈ –T_22__imaginary (i.e., 2.610226e+02 ≈ (–) – 2.605707e+02). The complex number (–a + ib) is a point T_21_ in the second quadrant. The complex number (c – id) is a point T_22_ in the fourth quadrant. The principal value of the argument (φ_0_) of a complex number (x + iy) is given by the following formulae:

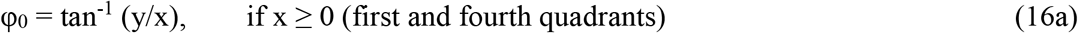

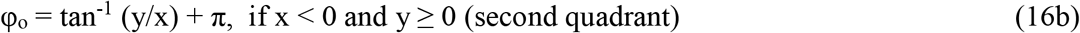

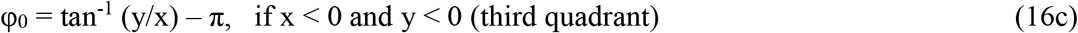

where the angle φ_0_ is in radian. (The angle in radian is given in degree (º) through: π radians = 180º. Therefore, 1 radian = 180º / 3.141593 ≈ 57.296º.) In (15a), the angle φ_0_ is: 144º. In (15b), the angle φ_0_ is: –36º. (In the rectangular system of coordinates, the positive angles are measured in the anticlockwise direction. Also, the negative angles are measured in the clockwise direction.) The expression of φ_0_ in (15a) describes electromagnetic waves, propagating in the microcavity, with a phase (angle) of 144°. Likewise, the expression of φ_0_ in (15b) describes electromagnetic waves with a phase (angle) of –36º. (Conventionally, the electromagnetic waves with phase (angle) negative and positive are treated as “outgoing” and “incoming” waves, respectively.) There is a phase change of 144º – (–36º) = 180º of electromagnetic waves propagating in the microcavity. The wavelength of resonance is λ = 1385 nm, at which the phase (angle) abruptly changes sign from 144º to –36º. (Here, φ_0_ becomes positive-to-negative. The points T_21_ and T_22_ are in the second and forth quadrants, respectively. Or, φ_0_ becomes negative– to–positive, when the points T_21_ and T_22_ are in the fourth and second quadrants, respectively. In an analogous manner, the condition of resonance is “high” (“h”) when the points T_21_ and T_22_ are in the first and third quadrants, respectively. The corresponding φ_0_ are positive and negative, respectively. Or, the points T_21_ and T_22_ are in the third and first quadrants, respectively. The corresponding φ_0_ are negative and positive, respectively. There is a phase change of 180º in each of these cases.) In physical realism, the electromagnetic waves undergo a number of rotations of 360º in the dielectric microcavity at the computed propagation (resonant) wavelength (λ), delivering excessive photon energy to the dielectric medium. The number of rotations in a microcavity, and hence the transfer of photon energy, may be enhanced by increasing the number of pairs of Bragg “reflector” layers (Section 6.1.1).

In the present computations, the set of calculations in the UVB band are performed to one-decimal-place in wavelength (δλ = 0.1 nm). The set of calculations in the UVA band are performed to one-place in wavelength (δλ = 1 nm). A range of wavelengths (λ) of 3 nm in the UVB band and 5 nm in the UVA band, where the idealized keratinocyte microcavity resonates, are considered to be the electromagnetic waves that propagate and transfer excessive photon energy in the epidermis. For the keratinocyte microcavity of varied diameters (15 µm, 30 µm, 50 µm and 70 µm), there is observed no wavelength that may reveal the condition of “high” (“h”) resonance of microcavity (T_21__real ≈ –T_22__real, T_21__imaginary ≈ –T_22__imaginary, phase change ≈ 180º) in the UVB and UVA bands.

### 6.1 Design and Analysis of a Spherical Dielectric Microcavity in the Epidermis of White-Skin in the UVB and UVA Bands

A spherical dielectric microcavity, representing an idealized keratinocyte in the epidermis of White-skin, consists of core of melanin (diameter = 800 nm, radius = 400 nm), and a pair of layers of keratinocyte ((i) diameter = 15 µm, radius = 7.5 µm, (ii) diameter = 30 µm, radius = 15 µm, (iii) diameter = 50 µm, radius = 25 µm, (iv) diameter = 70 µm, radius = 35 µm) and keratin (physical dimension not needed in the computations).

#### 6.1.1 Spherical Dielectric Microcavity for White-Skin Epidermis in the UVB Band

(i) Spherical dielectric microcavity in the UVB band, with keratinocyte of diameter of 15 µm, has the following parameters (Figure 1):

**Figure 1.**
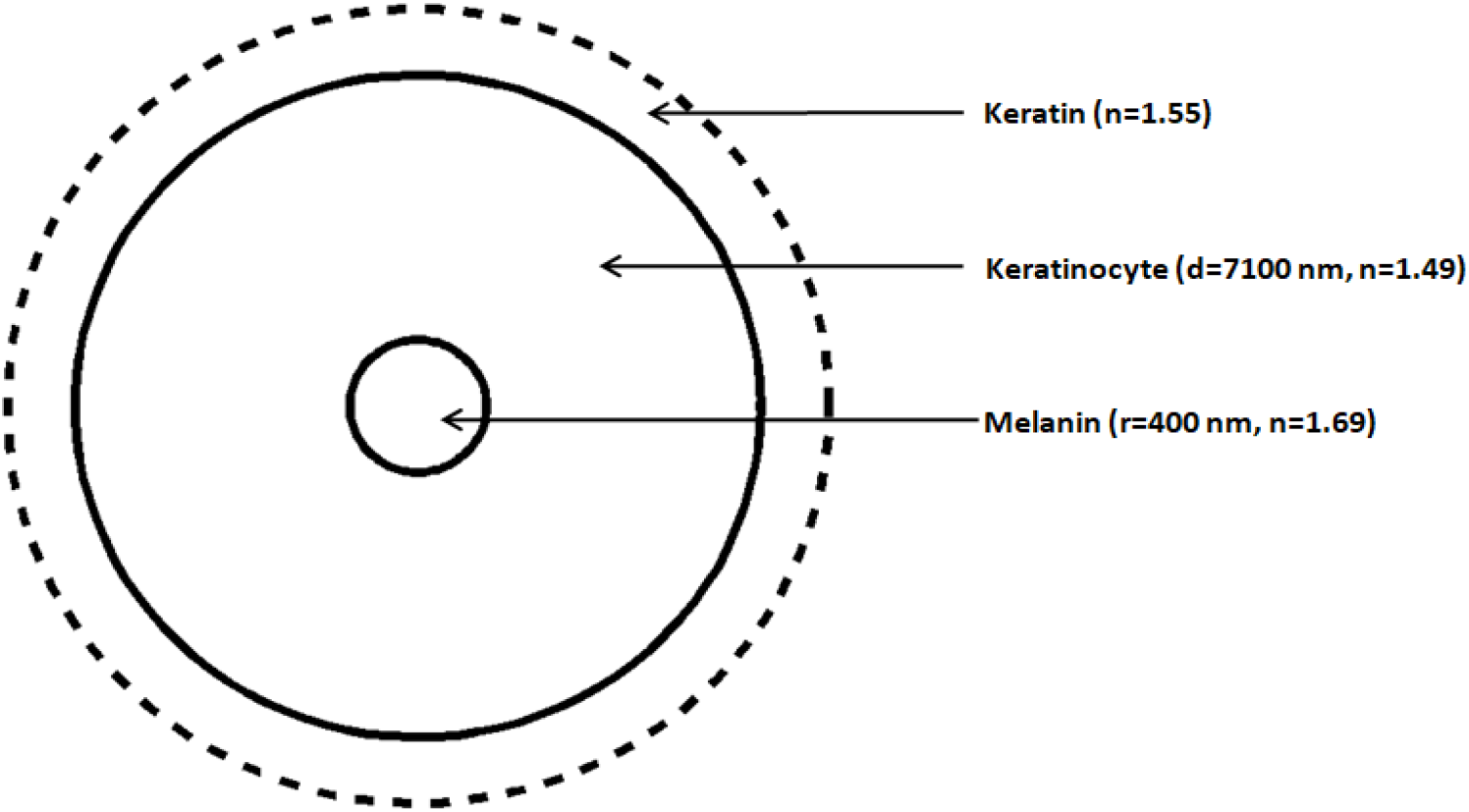
A spherical dielectric microcavity represents an idealized keratinocyte of diameter 15 µm in the epidermis of White-skin in the UVB band. It consists of core of melanin (radius r = 400 nm, refractive index n = 1.69) and a pair of layers of keratinocyte (physical dimension d = 7100 nm, refractive index n = 1.49) and keratin (dotted, refractive index n = 1.55).

**Figure 2.**
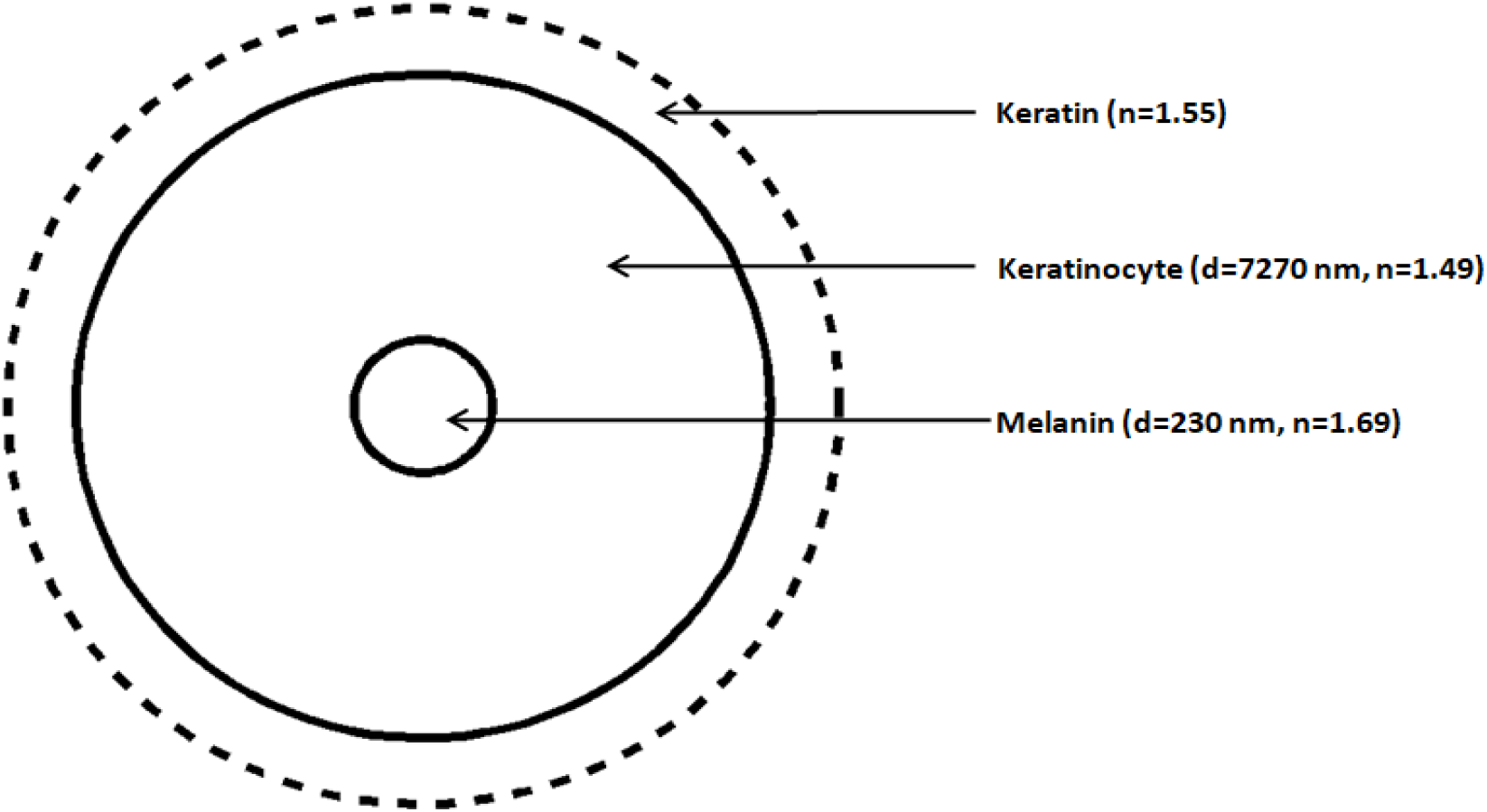
A spherical dielectric microcavity represents an idealized keratinocyte of diameter 15 µm in the epidermis of Pale Yellow-skin in the UVB band. It consists of core of melanin (radius r = 230 nm, refractive index n = 1.69) and a pair of layers of keratinocyte (physical dimension d = 7270 nm, refractive index n = 1.49) and keratin (dotted, refractive index n = 1.55).

Core of melanin, radius = 400 nm

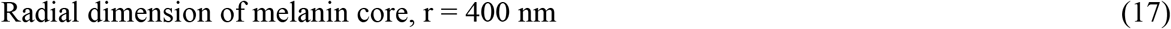

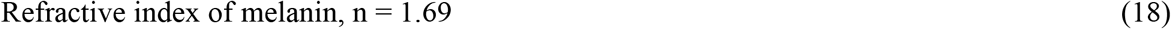

Layer of keratinocyte, radius = 7.5 µm = 7500 nm

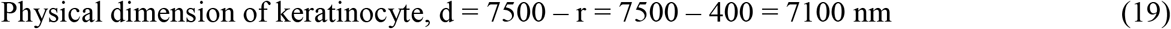

(From (17))

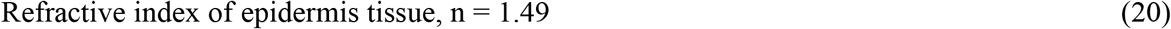

Layer of keratin (physical dimension not needed, dotted in Figure 1)

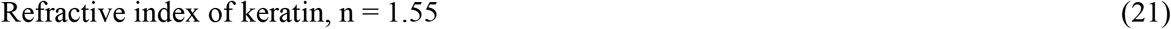

In a dielectric microcavity, the Bragg “reflection” wavelength λ_BR_ (nm) is given by

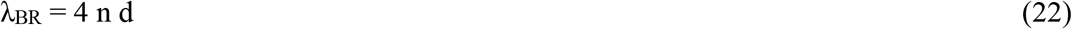

where d is the physical dimension (nm) of a dielectric layer of refractive index n. Here, the dielectric layer is the keratinocyte of dimension d = 7100 nm and refractive index n = 1.49, given by (19) and (20), respectively. The Bragg wavelength (λ_BR_, nm) is,

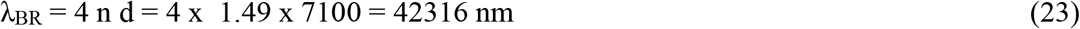

(Alternately, for a given (λ_BR_, nm), the physical dimension d (nm) of a dielectric layer of refractive index n is given by d = λ_BR_ / 4 n.) The Bragg wavelength λ_BR_ is the Bragg “reflection” wavelength. The keratinocyte layer is the Bragg “reflector” layer. The spherical microcavity is said to have been designed for the Bragg wavelength λ_BR_ = 42316 nm (≈ 0.042 mm) in the infrared band of the solar radiation spectrum (about 100 nm to 1 mm). The wavelength of propagation of electromagnetic waves λ equals λ_BR_ by suitably increasing the number of pairs of Bragg “reflector” layers.

For illustration, consider the aforementioned “designed” spherical microcavity consisting of core of ZnTe (r = 232 nm, n = 2.7) and a pair of layers of SiO_2_ (d = 267 nm, n = 1.45) and ZnTe (d = 144 nm, n = 2.7) (Section 6) [22]. The spherical microcavity is designed for the Bragg wavelength λ_BR_ = 1550 nm. The microcavity resonates at the wavelength λ = 1385 nm. Thus, the wavelength of propagation is computed as: λ = 1385 nm. However, with eight pairs of layers of ZnTe and SiO_2_, and radial dimension of core ZnTe of 2092 nm, λ = λ_BR_ = 1550 nm [22].

In the present computations, the complex matrix T (2 x 2) is calculated at δλ = 0.5 nm (and δλ = 0.1 nm for finer calculations) in the UVB band. The condition of occurrence of resonance of dielectric microcavity is examined at each wavelength λ, i.e., T_21__real ≈ –T_22__real and T_21__imaginary ≈ –T_22__imaginary. For the keratinocyte of diameter 15 µm, the microcavity resonates, with the point T_21_ (= –T_21__real + iT_21__imaginary) in the second quadrant (φ_0_ positive) and the point T_22_ (= T_22__real + iT_22__imaginary) in the first quadrant (φ_0_ positive). The condition of resonance of microcavity is deemed to be “good” (“g”) in the wavelength range 296.5 – 299.8 nm (T_21__real ≈ –T_22__real, T_21__imaginary ≠ –T_22__imaginary, phase change ≠ 180º, ratio of magnitudes of T_21__real and T_22__real ≤ 6). Also, the condition of resonance of microcavity is deemed to be “fair” (“f”) at the individual wavelengths of 296.4 nm and 299.9 nm (T_21__real ∼ – T_22__real, T_21__imaginary ≠ –T_22__imaginary, phase change ≠ 180º, ratio of magnitudes of T_21__real and T_22__real ≈ 6 to 10). The condition of resonance of microcavity is deemed to be “not met” at the wavelength of 300.0 nm, since the ratio of magnitudes of T_21__real and T_22__real ≈ 12.7. Alternately, it could be |T_22__real| / |T_21__real| ≤ 6, or 6 to 10, etcetera. The data below are the output of the computer program in C–programming language cited in Section 6; e is the base-10 of a number [24a, 24b, 24c]. The wavelengths of resonance of spherical dielectric microcavity are depicted in Table 6 (Section 6.1.3).

For illustration, the elements T_21_ and T_22_ of the complex matrix T (2 x 2) at the select wavelengths are the following:

a. λ = 296.4 nm, -2.184028e-02, 6.475932e-02 (T_21__real, T_21__imaginay) 2.098580e-03, 8.810022e-01 (T_22__real, T_22__imaginary) The condition of resonance is “fair” (“f”), with |T_21__real| / |T_22__real| ≈ 10.5.
b. λ = 298 nm, -1.493947e-02, 3.590073e-02 (T_21__real, T_21__imaginay) 3.802736e-02, 8.783873e-01 (T_22__real, T_22__imaginary) The condition of resonance is “good” (“g”), with |T_22__real| / |T_21__real| ≈ 2.5.
c. λ = 299.9 nm, -1.003193e-02, 7.249197e-02 (T_21__real, T_21__imaginay) 8.153281e-02, 8.776139e-01 (T_22__real, T_22__imaginary) The condition of resonance is “fair” (“f”), with |T_22__real| / |T_21__real| ≈ 8.2.
d. λ = 300.0 nm, -6.640119e-03, 7.305973e-02 (T_21__real, T_21__imaginay) 8.394770e-02, 8.774011e-01 (T_22__real, T_22__imaginary) The condition of resonance is deemed to be “not met”, since |T_22__real| / |T_21__real| ≈ 12.7.

(ii) Spherical dielectric microcavity in the UVB band, with keratinocyte of diameter of 30 µm, has the following parameters:

Core of melanin, radius = 400 nm

Radial dimension of melanin core, r = 400 nm

Refractive index of melanin, n = 1.69

Layer of keratinocyte, radius = 15 µm = 15000 nm

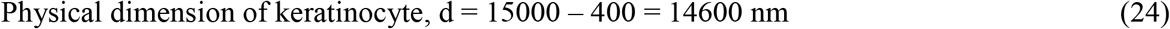

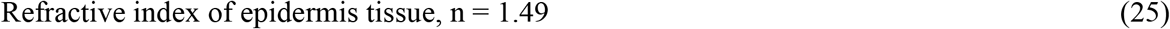

Layer of keratin (physical dimension not needed)

Refractive index of keratin, n = 1.55

The spherical microcavity is said to have been designed for the Bragg wavelength λ_BR_ = 4 x 1.49 x 14600 = 87016 nm (≈ 0.087 mm) in the infrared band of the solar radiation spectrum (from (24) and (25)).

For the keratinocyte of diameter 30 µm, the point T_21_ is in the first quadrant (φ_0_ positive) and the point T_22_ is in the fourth quadrant (φ_0_ negative). The condition of resonance of microcavity is “fair” (“f”) in the wavelength range 280.0 – 283.6 nm (T_21__imaginary ∼ – T_22__imaginary, T_21__real ≠ –T_22__real, phase change ≠ 180º, |T_21__imaginary| / |T_22__imaginary| ≈ 6 to 10). The wavelengths of resonance of spherical dielectric microcavity are depicted in Table 6 (Section 6.1.3).

(iii) Spherical dielectric microcavity in the UVB band, with keratinocyte of diameter of 50 µm, has the following parameters:

Core of melanin, radius = 400 nm

Radial dimension of melanin core, r = 400 nm

Refractive index of melanin, n = 1.69

Layer of keratinocyte, radius = 25 µm = 25000 nm

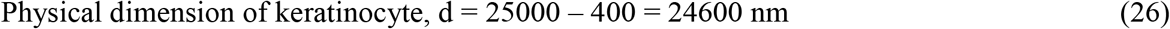

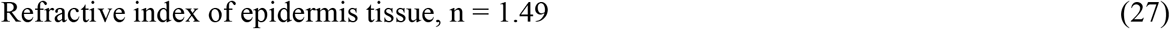

Layer of keratin (physical dimension not needed)

Refractive index of keratin, n = 1.55

The spherical microcavity is said to have been designed for the Bragg wavelength λ_BR_ = 4 x 1.49 x 24600 = 146616 nm (≈ 0.15 mm) in the infrared band of the solar radiation spectrum (from (26) and (27)).

For the keratinocyte of diameter 50 µm, the point T_21_ is in the first quadrant (φ_0_ positive) and the point T_22_ is in the fourth quadrant (φ_0_ negative). The condition of resonance of microcavity is “good” (“g”) in the wavelength range 284.2 – 287.1 nm (T_21__imaginary ≈ – T_22__imaginary, T_21__real ≠ –T_22__real, phase change ≠ 180º, |T_21__imaginary| / |T_22__ imaginary| ≤ 6). The wavelengths of resonance of spherical dielectric microcavity are depicted in Table 6 (Section 6.1.3).

(iv) Spherical dielectric microcavity in the UVB band, with keratinocyte of diameter of 70 µm, has the following parameters:

Core of melanin, radius = 400 nm

Radial dimension of melanin core, r = 400 nm

Refractive index of melanin, n = 1.69

Layer of keratinocyte, radius = 35 µm = 35000 nm

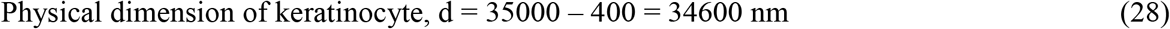

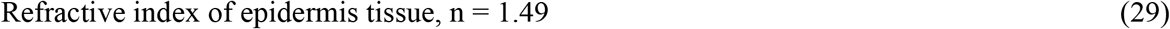

Layer of keratin (physical dimension not needed)

Refractive index of keratin, n = 1.55

The spherical microcavity is said to have been designed for the Bragg wavelength λ_BR_ = 4 x 1.49 x 34600 = 206216 nm (≈ 0.20 mm) in the infrared band of the solar radiation spectrum (from (28) and (29)).

For the keratinocyte of diameter 70 µm, the point T_21_ is in the second quadrant (φ_0_ positive) and the point T_22_ is in the first quadrant (φ_0_ positive). The condition of resonance of microcavity is “fair” (“f”) in the wavelength range 280.0 – 283.0 nm (T_21__real ∼ –T_22__real, T_21__imaginary ≠ –T_22__imaginary, phase change ≠ 180º, |T_21__real| / |T_22__real| ≈ 6 to 10). The wavelengths of resonance of microcavity are depicted in Table 6 (Section 6.1.3).

#### 6.1.2 Spherical Dielectric Microcavity for White-Skin Epidermis in the UVA Band

(i) Spherical dielectric microcavity in the UVA band, with keratinocyte of diameter of 15 µm, has the following parameters:

Core of melanin, radius = 400 nm

Radial dimension of melanin core, r = 400 nm

Refractive index of melanin, n = 1.67

Layer of keratinocyte, radius = 7.5 µm = 7500 nm

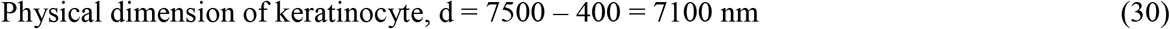

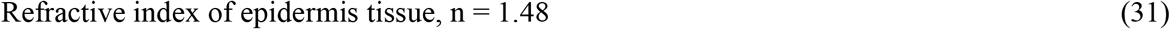

Layer of keratin (physical dimension not needed)

Refractive index of keratin, n = 1.55

The spherical microcavity is said to have been designed for the Bragg wavelength λ_BR_ = 4 x 1.48 x 7100 = 42032 nm (≈ 0.042 mm) in the infrared band of the solar radiation spectrum (from (30) and (31)).

The calculations in the UVA (315 – 400 nm) band are performed to integer-wavelength (δλ = 1 nm) in order to detect occurrence of resonance of dielectric microcavity. For the keratinocyte of diameter 15 µm, the point T_21_ is in the second quadrant (φ_0_ positive) and the point T_22_ is in the first quadrant (φ_0_ positive). The condition of resonance of microcavity is “good” (“g”) in the wavelength range 361 – 371 nm (T_21__real ≈ –T_22__real, T_21__imaginary ≠ – T_22__imaginary, phase change ≠ 180º, |T_21__real | / |T_22__real | ≤ 6). The wavelengths of resonance of spherical dielectric microcavity are depicted in Table 6 (Section 6.1.3).

(ii) Spherical dielectric microcavity in the UVA band, with keratinocyte of diameter of 30 µm, has the following parameters:

Core of melanin, radius = 400 nm

Radial dimension of melanin core, r = 400 nm

Refractive index of melanin, n = 1.67

Layer of keratinocyte, radius = 15 µm = 15000 nm

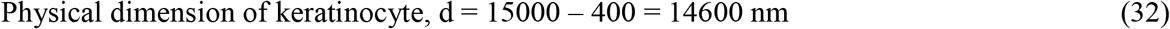

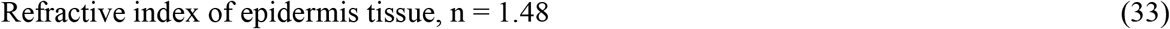

Layer of keratin (physical dimension not needed)

Refractive index of keratin, n = 1.55

The spherical microcavity is said to have been designed for the Bragg wavelength λ_BR_ = 4 x 1.48 x 14600 = 86432 nm (≈ 0.086 mm) in the infrared band of the solar radiation spectrum (from (32) and (33)).

For the keratinocyte of diameter 30 µm: (a) The point T_21_ is in the third quadrant (φ_0_ negative) and the point T_22_ is in the first quadrant (φ_0_ positive). The condition of resonance of microcavity is similar to that of “high” (“h”) resonance (Section 6). The condition of resonance of microcavity is “fair” (“f”) in the wavelength range 317 – 320 nm (T_21__imaginary ∼ – T_22__imaginary, T_21__real ≠ –T_22__real, phase change ≠ 180º, |T_21__imaginary| / |T_22__imaginary| ≈ 6 to 10). Also, the condition of resonance of microcavity is “good” (“g”) in the wavelength range 321 – 326 nm (T_21__imaginary ≈ –T_22__imaginary, T_21__real ≠ –T_22__real, phase change ≠ 180º, |T_21__imaginary| / |T_22__ imaginary| ≤ 6). (b) The point T_21_ is in the fourth quadrant (φ_0_ negative) and the point T_22_ is in the third quadrant (φ_0_ negative). The condition of resonance of microcavity is “good” (“g”) in the wavelength range 355 – 362 nm (T_21__real ≈ –T_22__real, T_21__imaginary ≠ –T_22__imaginary, phase change ≠ 180º, |T_21__real| / |T_22__real| ≤ 6). The wavelengths of resonance of spherical dielectric microcavity are depicted in Table 6 (Section 6.1.3).

(iii) Spherical dielectric microcavity in the UVA band, with keratinocyte of diameter of 50 µm, has the following parameters:

Core of melanin, radius = 400 nm

Radial dimension of melanin core, r = 400 nm

Refractive index of melanin, n = 1.67

Layer of keratinocyte, radius = 25 µm = 25000 nm

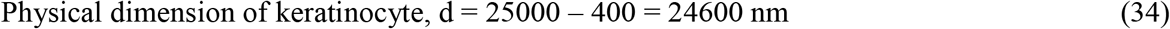

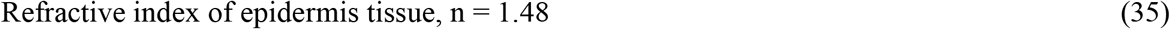

Layer of keratin (physical dimension not needed)

Refractive index of keratin, n = 1.55

The spherical microcavity is said to have been designed for the Bragg wavelength λ_BR_ = 4 x 1.48 x 24600 = 145632 (≈ 0.15 mm) in the infrared band of the solar radiation spectrum (from (34) and (35)).

For the keratinocyte of diameter 50 µm, the point T_21_ is in the fourth quadrant (φ_0_ negative) and the point T_22_ is in the third quadrant (φ_0_ negative). The condition of resonance of microcavity is “good” (“g”) in the wavelength range 353 – 356 nm (T_21__real ≈ –T_22__real, T_21__imaginary ≠ –T_22__imaginary, phase change ≠ 180º, |T_21__real| / |T_22__real| ≤ 6). Also, the condition of resonance of microcavity is “fair” (“f”) in the wavelength range 357 – 359 nm (T_21__real ∼ –T_22__real, T_21__imaginary ≠ –T_22__imaginary, phase change ≠ 180º, |T_21__real| / |T_22__real| ≈ 6 to 10). The wavelengths of resonance of spherical dielectric microcavity are depicted in Table 6 (Section 6.1.3).

(iv) Spherical dielectric microcavity in the UVA band, with keratinocyte of diameter of 70 µm, has the following parameters:

Core of melanin, radius = 400 nm

Radial dimension of melanin core, r = 400 nm

Refractive index of melanin, n = 1.67

Layer of keratinocyte, radius = 35 µm = 35000 nm

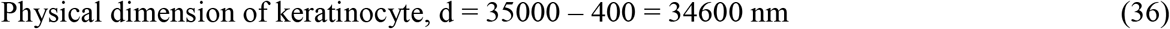

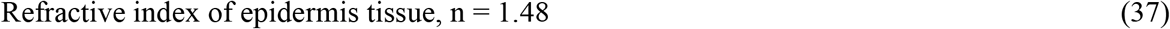

Layer of keratin (physical dimension not needed)

Refractive index of keratin, n = 1.55

The spherical microcavity is said to have been designed for the Bragg wavelength λ_BR_ = 4 x 1.48 x 34600 = 204832 (≈ 0.20 mm) in the infrared band of the solar radiation spectrum (from (36) and (37)).

For the keratinocyte of diameter 70 µm, the point T_21_ is in the fourth quadrant (φ_0_ negative) and the point T_22_ is in the third quadrant (φ_0_ negative). The condition of resonance of microcavity is “good” (“g”) in the wavelength range 352 – 355 nm (T_21__real ≈ –T_22__real, T_21__imaginary ≠ –T_22__imaginary, phase change ≠ 180º, |T_21__real| / |T_22__real| ≤ 6). Also, the condition of resonance of microcavity is “fair” (“f”) in the wavelength range 356 – 357 nm (T_21__real ∼ –T_22__real, T_21__imaginary ≠ –T_22__imaginary, phase change ≠ 180º, |T_21__real| / |T_22__real| ≈ 6 to 10). The wavelengths of resonance of spherical dielectric microcavity are depicted in Table 6 (Section 6.1.3).

#### 6.1.3 A Summary and Analysis of Wavelengths of Resonance of Spherical Dielectric Microcavity for White-Skin Epidermis in the UVB and UVA Bands

The wavelengths of resonance of spherical dielectric microcavity for White-skin epidermis in the UVB and UVA bands are given in Table 6.

The propagation of electromagnetic waves in the UVB and UVA bands in the spherical dielectric microcavity in the epidermis of White-skin enables to determine the efficacy of the model study. The resonance range of wavelengths of 3 nm and 5 nm, under the approximation of “fair” (“f”) / “good” (“g”) condition of resonance, for the UVB and UVA bands, respectively, is thus fixed. The range of wavelengths of resonance is the model (model computed) propagation range of wavelengths, where photons are absorbed in the epidermis cells keratinocytes.

In the UVB band, in Table 6, the photons of wavelengths 296.4 nm to 299.9 nm are absorbed in the epidermis of White-skin. The corresponding keratinocytes of size 15 µm refer to the epidermis close to the basal layer, and the dermis. Thus the UVB photons of wavelengths 296.4 nm to 299.9 nm are absorbed in the epidermis close to the dermis. The model propagation range of wavelengths 296.4 nm (≈ 296 nm) to 299.9 nm (≈ 300 nm) is expectedly greater than 290 nm for the solar UVB radiation to be effective on the human skin. Also, the model propagation range closely matches the maximal range 295 nm to 300 nm, of the overall wavelength range 290 nm to ∼317 nm, of the absorption of UVB photons by 7-dehydrocholesterol in the human skin to synthesize vitamin D_3_ [1]. (The peak absorption by 7-dehydrocholesterol in the human skin is around 298 nm.) The model propagation range of 296 nm to 300 nm implies the effectiveness of the solar UVB radiation in the wavelength range 296 nm to 300 nm for the synthesis of vitamin D_3_. The synthesized vitamin D_3_ is transported to the body through the dermis. The keratinocytes of size 30 µm and 50 µm refer to the middle and the upper portions of the epidermis, respectively. The UVB photons of wavelengths 280.0 nm to 283.6 nm (≈ 284 nm) and 284.2 nm (≈ 284 nm) to 287.1 nm (≈ 287 nm) are absorbed in the middle and the upper portions of the epidermis, respectively. In the literature the keratinocytes of size 50 µm refer to the stratum corneum, where these are eliminated as corneocytes. The theoretically deduced keratinocytes of size 70 µm refer to the epidermis close to the stratum corneum. The UVB photons of wavelengths 280.0 nm to 283.0 nm are absorbed in the epidermis close to the stratum corneum. The model propagation range of 280 nm to 287.1 nm (≈ 287 nm) implies the application of UVB radiation phototherapies in the wavelength range 280 nm to 287, besides in the wavelength range 296 nm to 300 nm, toward synthesis of vitamin D_3_ in White-skin (Section 7). The model propagation range of 296 nm to 300 nm implies the causative effect of the excessive exposure of solar UVB radiation in the wavelength range 296 nm to 300 nm of “delayed” long-term tanning of White-skin.

In the UVA band, in Table 6, the photons of wavelengths 352 nm to 371 nm are absorbed in the epidermis of White-skin. The model propagation range closely matches the peak range of tanning of skin of 340 nm to 370 nm [7, Review]. The model propagation range implies the causative effect of the solar UVA radiation in the wavelength range 352 nm to 371 nm of “immediate” tanning of White-skin. Also, the model propagation range implies the application of UVA radiation phototherapies toward “immediate” tanning of White-skin (Section 7). The UVA photons of a narrow range of wavelengths 317 nm to 326 nm are absorbed in the keratinocytes of size 30 µm, that is, in the middle portion of the epidermis. The latter range of propagation wavelengths is due to the underlying “fair”/ “good” condition of resonance of spherical dielectric microcavity in the model study, and is summarily rejected. (The theoretical formulation of the spherical dielectric microcavity may result in single wavelength of resonance under the condition of “high” (“h”) resonance.)

### 6.2 Design and Analysis of a Spherical Dielectric Microcavity in the Epidermis of Pale Yellow-Skin in the UVB and UVA Bands

A spherical dielectric microcavity, representing an idealized keratinocyte in the epidermis of Pale Yellow-skin, consists of core of melanin (diameter = 460 nm, radius = 230 nm), and a pair of layers of keratinocyte ((i) diameter = 15 µm, radius = 7.5 µm, (ii) diameter = 30 µm, radius = 15 µm, (iii) diameter = 50 µm. radius = 25 µm, (iv) diameter = 70 µm, radius = 35 µm) and keratin (physical dimension not needed in the computations).

#### 6.2.1 Spherical Dielectric Microcavity for Pale Yellow-Skin Epidermis in the UVB Band

(i) Spherical dielectric microcavity in the UVB band, with keratinocyte of diameter of 15 µm, has the following parameters (Figure 2):

Core of melanin, radius = 230 nm

Radial dimension of melanin core, r = 230 nm

Refractive index of melanin, n = 1.69

Layer of keratinocyte, radius = 7.5 µm = 7500 nm

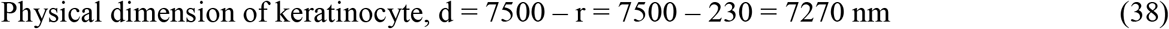

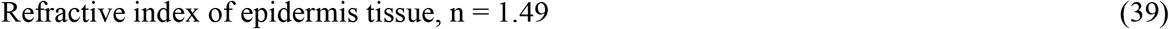

Layer of keratin (physical dimension not needed, dotted in Figure 2)

Refractive index of keratin, n = 1.55

The spherical microcavity is said to have been designed for the Bragg wavelength λ_BR_ = 4 x 1.49 x 7270 = 43329 nm (≈ 0.043 mm) in the infrared band of the solar radiation spectrum (from (38) and (39)).

For the keratinocyte of diameter 15 µm, the condition of resonance of microcavity is “good” (“g”) in the wavelength range 282.0 – 283.5 nm. Also, the condition of resonance of microcavity is “fair” (“f”) in the wavelength range 283.6 – 287.3 nm. The wavelengths of resonance of spherical dielectric microcavity are depicted in Table 7 (Section 6.2.3).

**Table 7.** The wavelengths of resonance of spherical dielectric microcavity representing melanin embedded keratinocyte of size 15 µm to 70 µm in the epidermis of Pale Yellow-skin in the UVB and UVA bands. The condition of resonance is termed as “good” (“g”) or “fair” (“f”). A range of 3 nm in the UVB and 5 nm in the UVA is considered as the wavelengths of resonance. A spherical dielectric microcavity not resonating at a keratinocyte size is represented by “–”

| Wavelength Bands ($\lambda$) | Pale yellow-skin Keratinocyte Size $15 \text{ }\mu\text{m}$ | Pale yellow-skin Keratinocyte Size $30 \text{ }\mu\text{m}$ | Pale yellow-skin Keratinocyte Size $50 \text{ }\mu\text{m}$ | Pale yellow-skin Keratinocyte Size $70 \text{ }\mu\text{m}$ |
| --- | --- | --- | --- | --- |
| UVB<br>(280 – 315 nm) | 282.0–283.5 nm (“g”) | 285.1–290.7 nm (“g”) | 291.1–293.6 nm (“g”) | – |
|  | –<br>283.6–287.3 nm (“f”) | –<br>290.8–292.5 nm (“f”) | –<br>293.7–294.7 nm (“f”) |  |
| UVA<br>(315 – 400 nm) | 360–371 nm (“f”) | 330–332 nm (“f”) | 352–354 nm (“f”) | 351–352 nm (“f”) |
|  | –<br>372–375 nm (“g”) | –<br>333–336 nm (“g”) | –<br>355–359 nm (“g”) | –<br>353–356 nm (“g”) |
|  |  | 359–367 nm (“g”) | 375–380 nm (“g”) |  |

(ii) Spherical dielectric microcavity in the UVB band, with keratinocyte of diameter of 30 µm, has the following parameters:

Core of melanin, radius = 230 nm

Radial dimension of melanin core, r = 230 nm

Refractive index of melanin, n = 1.69

Layer of keratinocyte, radius = 15 µm = 15000 nm

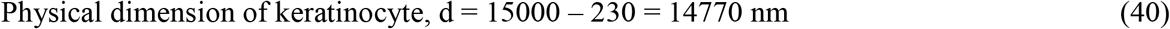

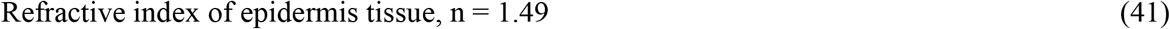

Layer of keratin (physical dimension not needed)

Refractive index of keratin, n = 1.55

The spherical microcavity is said to have been designed for the Bragg wavelength λ_BR_ = 4 x 1.49 x 14770 = 88029 nm (≈ 0.088 mm) in the infrared band of the solar radiation spectrum (from (40) and (41)).

For the keratinocyte of diameter 30 µm, the condition of resonance of microcavity is “good” (“g”) in the wavelength range 285.1 – 290.7 nm. Also, the condition of resonance of microcavity is “fair” (“f”) in the wavelength range 290.8 – 292.5 nm. The wavelengths of resonance of spherical dielectric microcavity are depicted in Table 7 (Section 6.2.3).

(iii) Spherical dielectric microcavity in the UVB band, with keratinocyte of diameter of 50 µm, has the following parameters:

Core of melanin, radius = 230 nm

Radial dimension of melanin core, r = 230 nm

Refractive index of melanin, n = 1.69

Layer of keratinocyte, radius = 25 µm = 25000 nm

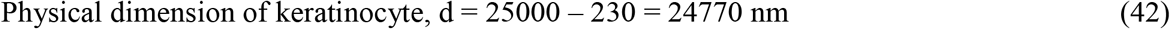

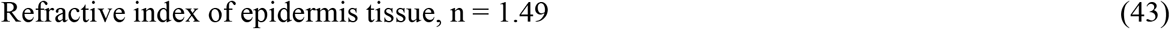

Layer of keratin (physical dimension not needed)

Refractive index of keratin, n = 1.55

The spherical microcavity is said to have been designed for the Bragg wavelength λ_BR_ = 4 x 1.49 x 24770 = 147629 nm (≈ 0.15 mm) in the infrared band of the solar radiation spectrum (from (42) and (43)).

For the keratinocyte of diameter 50 µm, the condition of resonance of microcavity is “good” (“g”) in the wavelength range 291.1 – 293.6 nm. Also, the condition of resonance of microcavity is “fair” (“f”) in the wavelength range 293.7 – 294.7 nm. The wavelengths of resonance of spherical dielectric microcavity are depicted in Table 7 (Section 6.2.3).

(iv) Spherical dielectric microcavity in the UVB band, with keratinocyte of diameter of 70 µm, has the following parameters:

Core of melanin, radius = 230 nm

Radial dimension of melanin core, r = 230 nm

Refractive index of melanin, n = 1.69

Layer of keratinocyte, radius = 35 µm = 35000 nm

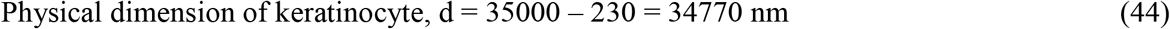

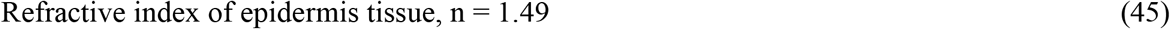

Layer of keratin (physical dimension not needed)

Refractive index of keratin, n = 1.55

The spherical microcavity is said to have been designed for the Bragg wavelength λ_BR_ = 4 x 1.49 x 34770 = 207229 nm (≈ 0.21 mm) in the infrared band of the solar radiation spectrum (from (44) and (45)).

For the keratinocyte of diameter 70 µm, the spherical dielectric microcavity does not reveal any resonance of the spherical dielectric microcavity (“–” in Table 7, Section 6.2.3).

#### 6.2.2 Spherical Dielectric Microcavity for Pale Yellow-Skin Epidermis in the UVA Band

(i) Spherical dielectric microcavity in the UVA band, with keratinocyte of diameter of 15 µm, has the following parameters:

Core of melanin, radius = 230 nm

Radial dimension of melanin core, r = 230 nm

Refractive index of melanin, n = 1.67

Layer of keratinocyte, radius = 7.5 µm = 7500 nm

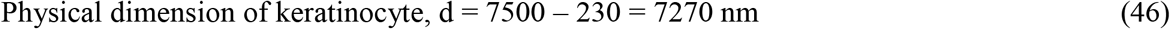

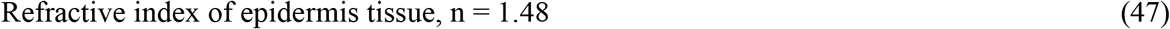

Layer of keratin (physical dimension not needed)

Refractive index of keratin, n = 1.55

The spherical microcavity is said to have been designed for the Bragg wavelength λ_BR_ = 4 x 1.48 x 7100 = 43038 nm (≈ 0.043 mm) in the infrared band of the solar radiation spectrum (from (46) and (47)).

For the keratinocyte of diameter 15 µm, the condition of resonance of microcavity is similar to that of “high” (“h”) resonance (Section 6). The condition of resonance of microcavity is “fair” (“f”) in the wavelength range 360 – 371 nm. Also, the condition of resonance of microcavity is “good” (“g”) in the wavelength range 372 – 375 nm. The wavelengths of resonance of spherical dielectric microcavity are depicted in Table 7 (Section 6.2.3).

(ii) Spherical dielectric microcavity in the UVA band, with keratinocyte of diameter of 30 µm, has the following parameters:

Core of melanin, radius = 230 nm

Radial dimension of melanin core, r = 230 nm

Refractive index of melanin, n = 1.67

Layer of keratinocyte, radius = 15 µm = 15000 nm

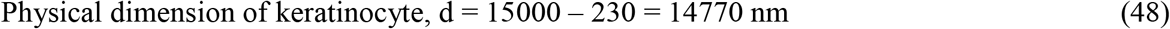

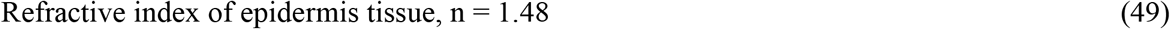

Layer of keratin (physical dimension not needed)

Refractive index of keratin, n = 1.55

The spherical microcavity is said to have been designed for the Bragg wavelength λ_BR_ = 4 x 1.48 x 14770 = 87438 nm (≈ 0.087 mm) in the infrared band of the solar radiation spectrum (from (48) and (49)).

For the keratinocyte of diameter 30 µm: (a) The condition of resonance of microcavity is “fair” (“f”) in the wavelength range 330 – 332 nm. Also, the condition of resonance of microcavity is “good” (“g”) in the wavelength range 333 – 336 nm. (b) The condition of resonance of microcavity is similar to that of “high” (“h”) resonance (Section 6). The condition of resonance of microcavity is “good” (“g”) in the wavelength range 359 – 367 nm. The wavelengths of resonance of spherical dielectric microcavity are depicted in Table 7 (Section 6.2.3).

(iii) Spherical dielectric microcavity in the UVA band, with keratinocyte of diameter of 50 µm, has the following parameters:

Core of melanin, radius = 230 nm

Radial dimension of melanin core, r = 230 nm

Refractive index of melanin, n = 1.67

Layer of keratinocyte, radius = 15 µm = 25000 nm

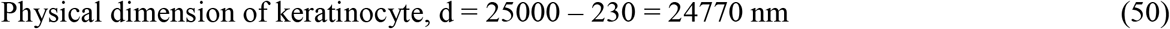

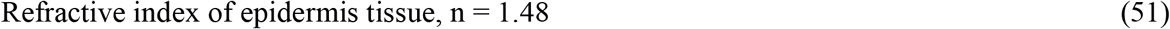

Layer of keratin (physical dimension not needed)

Refractive index of keratin, n = 1.55

The spherical microcavity is said to have been designed for the Bragg wavelength λ_BR_ = 4 x 1.48 x 24770 = 146638 nm (≈ 0.15 mm) in the infrared band of the solar radiation spectrum (from (50) and (51)).

For the keratinocyte of diameter 50 µm: (a) The condition of resonance of microcavity is “fair” (“f”) in the wavelength range 352 – 354 nm. Also, the condition of resonance of microcavity is “good” (“g”) in the wavelength range 355 – 359 nm. (b) The condition of resonance of microcavity is similar to that of “high” (“h”) resonance (Section 6). The condition of resonance of microcavity is “good” (“g”) in the wavelength range 375 – 380 nm. The wavelengths of resonance of spherical dielectric microcavity are depicted in Table 7 (Section 6.2.3).

(iv) Spherical dielectric microcavity in the UVA band, with keratinocyte of diameter of 70 µm, has the following parameters:

Core of melanin, radius = 230 nm

Radial dimension of melanin core, r = 230 nm

Refractive index of melanin, n = 1.67

Layer of keratinocyte, radius = 15 µm = 35000 nm

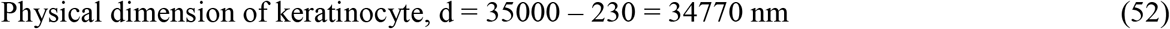

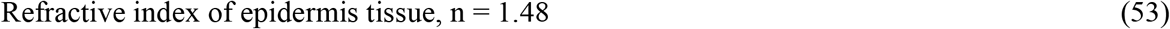

Layer of keratin (physical dimension not needed)

Refractive index of keratin, n = 1.55

The spherical microcavity is said to have been designed for the Bragg wavelength λ_BR_ = 4 x 1.48 x 34770 = 205838 nm (≈ 0.21 mm) in the infrared band of the solar radiation spectrum (from (52) and (53)).

For the keratinocyte of diameter 70 µm, the condition of resonance of microcavity is “fair” (“f”) in the wavelength range 351 – 352 nm. Also, the condition of resonance of microcavity is “good” (“g”) in the wavelength range 353 – 356 nm. The wavelengths of resonance of spherical dielectric microcavity are depicted in Table 7 (Section 6.2.3).

#### 6.2.3 A Summary and Analysis of Wavelengths of Resonance of Spherical Dielectric Microcavity for Pale Yellow-Skin Epidermis in the UVB and UVA Bands

The wavelengths of resonance of spherical dielectric microcavity for Pale Yellow-skin epidermis in the UVB and UVA bands are given in Table 7.

In the UVB band, in Table 7, the photons of wavelengths 282.0 nm to 294.7 nm are absorbed in the epidermis of Pale Yellow-skin. The model propagation range of wavelengths is partly greater than 290 nm for the solar UVB radiation to be effective on the human skin. Also, the model propagation range is amidst the wavelength range of 290 nm to ∼317 nm of the absorption of UVB – UVA photons by 7-dehydrocholesterol in the human skin to synthesize vitamin D_3_ [1]. The model propagation range implies the effectiveness of the solar UVB radiation in the wavelength range 290 nm to 294.7 nm (≈ 295 nm) for the synthesis of vitamin D_3_ in Pale Yellow-skin. The model propagation range also implies the application of UVB radiation therapies in the wavelength range 282 nm to 295 nm toward synthesis of vitamin D_3_ in Pale Yellow-skin (Section 7). The model propagation range of 290 nm to 295 nm implies the causative effect of the excessive exposure of solar UVB radiation in the wavelength range 290 nm to 295 nm of “delayed” long-term tanning of Pale Yellow-skin.

In the UVA band, in Table 7, the photons of wavelengths 351 nm to 380 nm are absorbed in the epidermis of Pale Yellow-skin. The model propagation range closely matches the peak range of tanning of skin of 340 nm to 370 nm [7, Review]. The model propagation range implies the causative effect of the solar UVA radiation in the wavelength range 351 nm to 380 nm of “immediate” tanning of Pale Yellow-skin. Also, the model propagation range implies the application of UVA radiation phototherapies toward “immediate” tanning of Pale Yellow-skin (Section 7). The UVA photons of wavelengths 330 nm to 336 nm are absorbed in the middle portion of the epidermis. The latter narrow range of propagation wavelengths is due to the underlying approximation of “fair”/ “good” condition of resonance of the spherical dielectric microcavity in the model study, and is summarily rejected.

### 6.3 Design and Analysis of a Spherical Dielectric Microcavity in the Epidermis of Brown-Skin in the UVB and UVA Bands

A spherical dielectric microcavity, representing an idealized keratinocyte in the epidermis of Brown-skin, consists of core of melanin (diameter = 320 nm, radius = 160 nm), and a pair of layers of keratinocyte ((i) diameter = 15 µm, radius = 7.5 µm, (ii) diameter = 30 µm, radius = 15 µm, (iii) diameter = 50 µm, radius = 25 µm, (iv) diameter = 70 µm, radius = 35 µm) and keratin (physical dimension not needed in the computations).

#### 6.3.1 Spherical Dielectric Microcavity for Brown-Skin Epidermis in the UVB Band

Spherical dielectric microcavity in the UVB band, with keratinocyte of diameter of 15 µm, has the following parameters (Figure 3):

**Figure 3.**
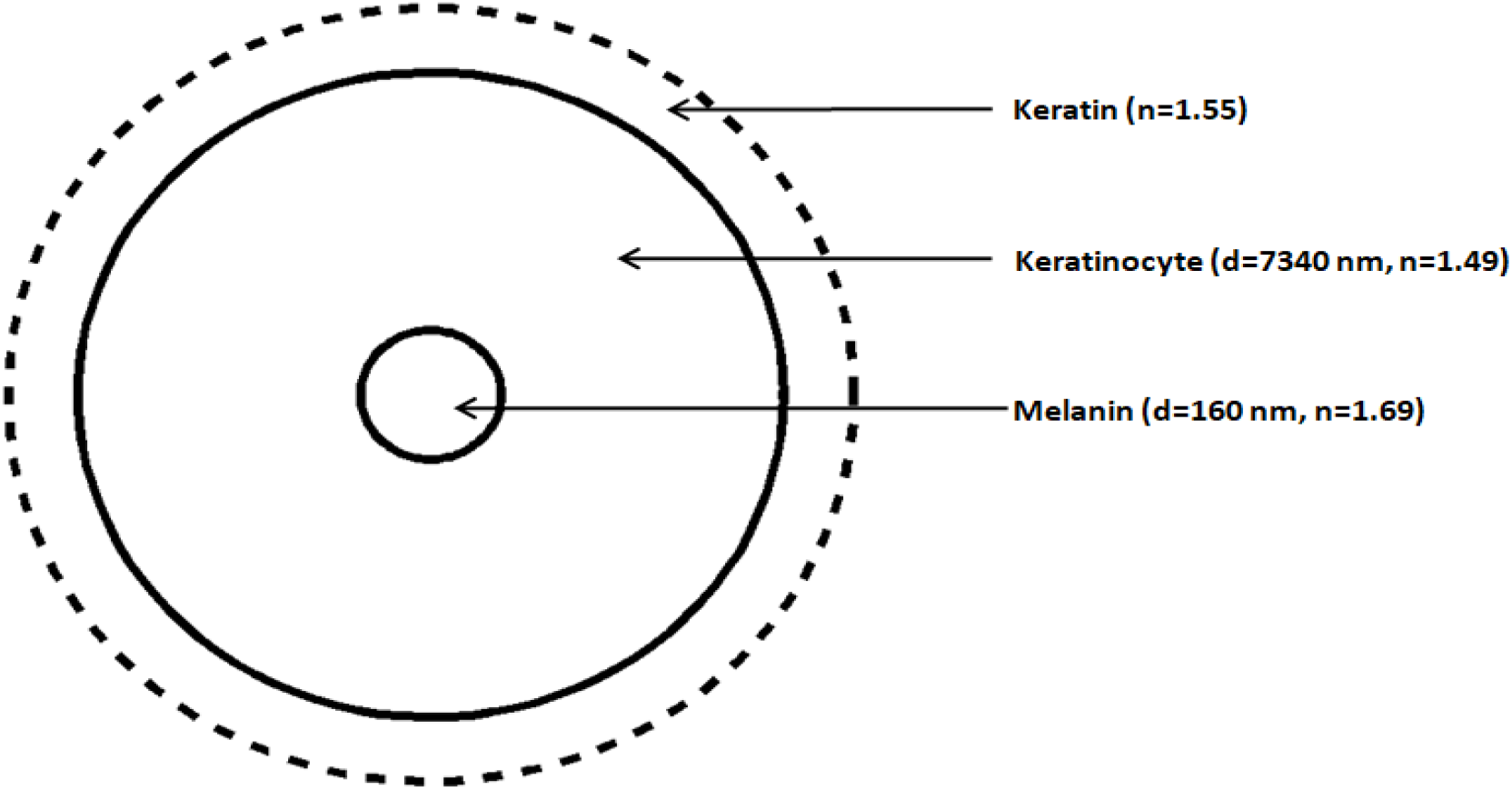
A spherical dielectric microcavity represents an idealized keratinocyte of diameter 15 µm in the epidermis of Brown-skin in the UVB band. It consists of core of melanin (radius r = 160 nm, refractive index n = 1.69) and a pair of layers of keratinocyte (physical dimension d = 7340 nm, refractive index n = 1.49) and keratin (dotted, refractive index n = 1.55).

Core of melanin, radius = 160 nm

Radial dimension of melanin core, r = 160 nm

Refractive index of melanin, n = 1.69

Layer of keratinocyte, radius = 7.5 µm = 7500 nm

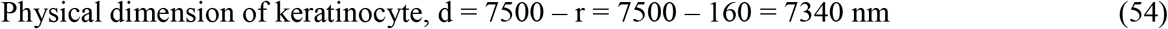

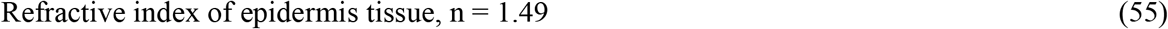

Layer of keratin (physical dimension not needed, dotted in Figure 3)

Refractive index of keratin, n = 1.55

The spherical microcavity is said to have been designed for the Bragg wavelength λ_BR_ = 4 x 1.49 x 7340 = 43746 nm (≈ 0.044 mm) in the infrared band of the solar radiation spectrum (from (54) and (55)).

For the keratinocyte of diameter 15 µm, the spherical dielectric microcavity does not reveal any resonance of the spherical dielectric microcavity (“–” in Table 8, Section 6.3.3).

**Table 8.**
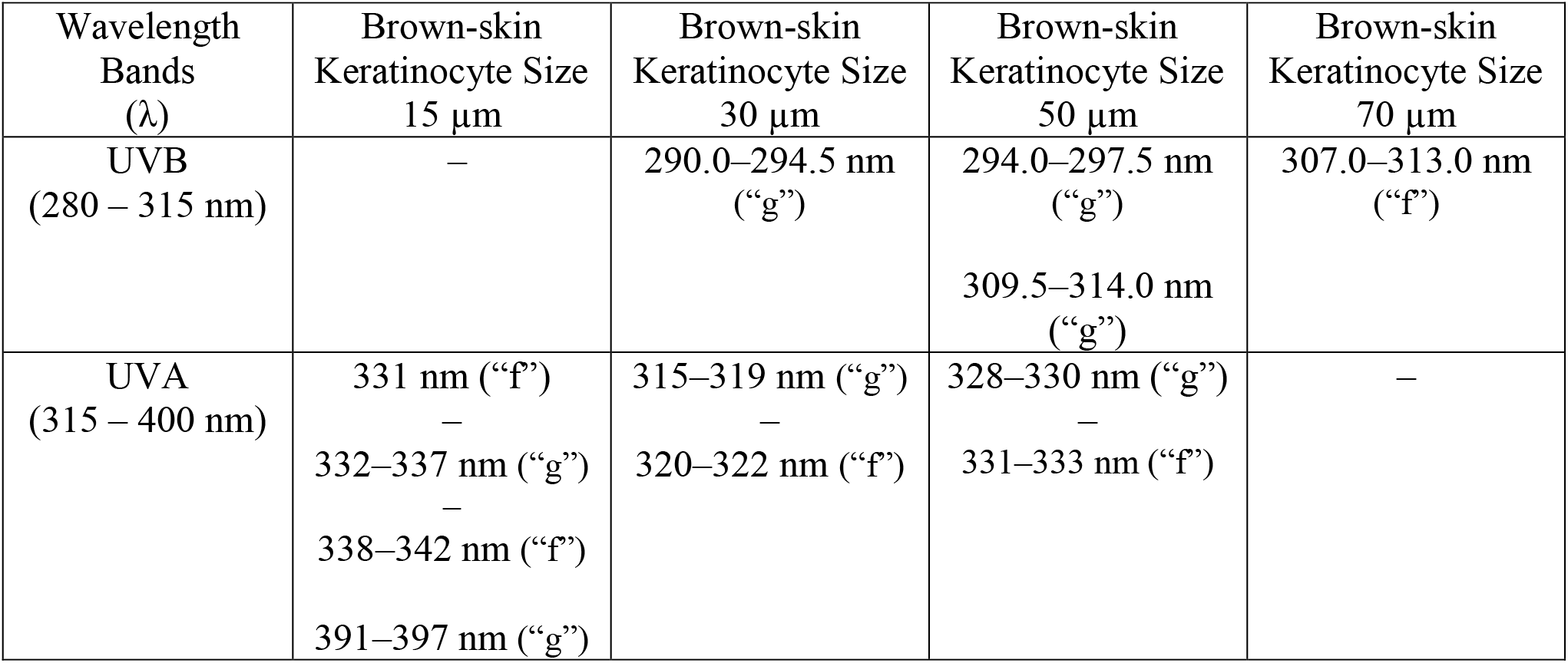
The wavelengths of resonance of spherical dielectric microcavity representing melanin embedded keratinocyte of size 15 µm to 70 µm in the epidermis of Brown-skin in the UVB and UVA bands. The condition of resonance is termed as “good” (“g”) or “fair” (“f”). A range of 3 nm in the UVB and 5 nm in the UVA is considered as the wavelengths of resonance. A spherical dielectric microcavity not resonating at a keratinocyte size is represented by “–”

Spherical dielectric microcavity in the UVB band, with keratinocyte of diameter of 30 µm, has the following parameters:

Core of melanin, radius = 160 nm

Radial dimension of melanin core, r = 160 nm

Refractive index of melanin, n = 1.69

Layer of keratinocyte, radius = 15 µm = 15000 nm

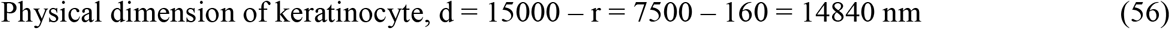

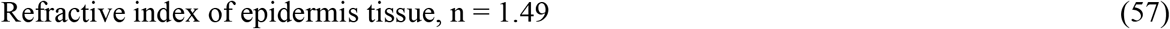

Layer of keratin (physical dimension not needed)

Refractive index of keratin, n = 1.55

The spherical microcavity is said to have been designed for the Bragg wavelength λ_BR_ = 4 x 1.49 x 14840 = 88446 nm (≈ 0.088 mm) in the infrared band of the solar radiation spectrum (from (56) and (57)).

For the keratinocyte of diameter 30 µm, the condition of resonance of microcavity is similar to that of “high” (“h”) resonance (Section 6). The condition of resonance of microcavity is “good” (“g”) in the wavelength range 290.0 – 294.5 nm. The wavelengths of resonance of spherical dielectric microcavity are depicted in Table 8 (Section 6.3.3).

Spherical dielectric microcavity in the UVB band, with keratinocyte of diameter of 50 µm, has the following parameters:

Core of melanin, radius = 160 nm

Radial dimension of melanin core, r = 160 nm

Refractive index of melanin, n = 1.69

Layer of keratinocyte, radius = 25 µm = 25000 nm

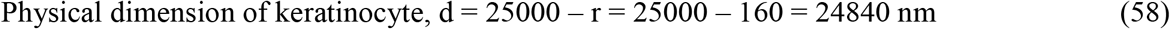

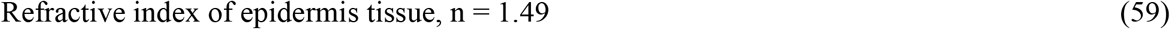

Layer of keratin (physical dimension not needed)

Refractive index of keratin, n = 1.55

The spherical microcavity is said to have been designed for the Bragg wavelength λ_BR_ = 4 x 1.49 x 24840 = 148046 nm (≈ 0.15 mm) in the infrared band of the solar radiation spectrum (from (58) and (59)).

For the keratinocyte of diameter 50 µm: (a) The condition of resonance of microcavity is similar to that of “high” (“h”) resonance (Section 6). The condition of resonance of microcavity is “good” (“g”) in the wavelength range 294.0 – 297.5 nm. (b) The condition of resonance of microcavity is “good” (“g”) in the wavelength range 309.5 – 314.0 nm. The wavelengths of resonance of spherical dielectric microcavity are depicted in Table 8 (Section 6.3.3).

Spherical dielectric microcavity in the UVB band, with keratinocyte of diameter of 70 µm, has the following parameters:

Core of melanin, radius = 160 nm

Radial dimension of melanin core, r = 160 nm

Refractive index of melanin, n = 1.69

Layer of keratinocyte, radius = 35 µm = 35000 nm

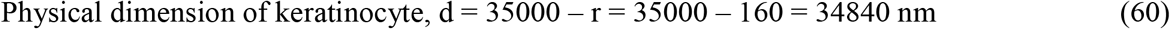

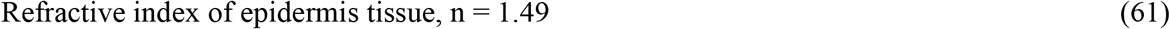

Layer of keratin (physical dimension not needed)

Refractive index of keratin, n = 1.55

The spherical microcavity is said to have been designed for the Bragg wavelength λ_BR_ = 4 x 1.49 x 34840 = 207646 nm (≈ 0.21 mm) in the infrared band of the solar radiation spectrum (from (60) and (61)).

For the keratinocyte of diameter 70 µm, the condition of resonance of microcavity is “fair” (“f”) in the wavelength range 307.0 – 313.0 nm. The wavelengths of resonance of spherical dielectric microcavity are depicted in Table 8 (Section 6.3.3).

#### 6.3.2 Spherical Dielectric Microcavity for Brown-Skin Epidermis in the UVA Band

(i) Spherical dielectric microcavity in the UVA band, with keratinocyte of diameter of 15 µm, has the following parameters:

Core of melanin, radius = 160 nm

Radial dimension of melanin core, r = 160 nm

Refractive index of melanin, n = 1.67

Layer of keratinocyte, radius = 7.5 µm = 7500 nm

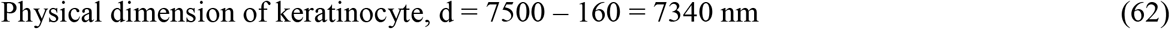

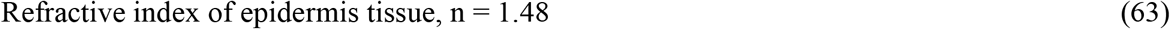

Layer of keratin (physical dimension not needed)

Refractive index of keratin, n = 1.55

The spherical microcavity is said to have been designed for the Bragg wavelength λ_BR_ = 4 x 1.48 x 7340 = 43453 nm (≈ 0.043 mm) in the infrared band of the solar radiation spectrum (from (62) and (63)).

For the keratinocyte of diameter 15 µm: (a) The condition of resonance of microcavity is “fair” (“f”) at the wavelength 331 nm. Also, the condition of resonance of microcavity is “good” (“g”) in the wavelength range 332 – 337 nm. Further, the condition of resonance of microcavity is “fair” (“f”) in the wavelength range 338 – 342 nm. (b) The condition of resonance of microcavity is “good” (“g”) in the wavelength range 391 – 397 nm. The wavelengths of resonance of spherical dielectric microcavity are depicted in Table 8 (Section 6.3.3).

(ii) Spherical dielectric microcavity in the UVA band, with keratinocyte of diameter of 30 µm, has the following parameters:

Core of melanin, radius = 160 nm

Radial dimension of melanin core, r = 160 nm

Refractive index of melanin, n = 1.67

Layer of keratinocyte, radius = 15 µm = 15000 nm

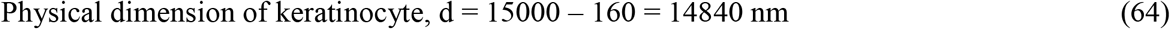

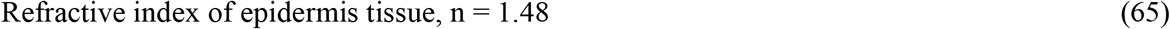

Layer of keratin (physical dimension not needed)

Refractive index of keratin, n = 1.55

The spherical microcavity is said to have been designed for the Bragg wavelength λ_BR_ = 4 x 1.48 x 14840 = 87853 nm (≈ 0.088 mm) in the infrared band of the solar radiation spectrum (from (64) and (65)).

For the keratinocyte of diameter 30 µm, the condition of resonance of microcavity is “good” (“g”) in the wavelength range 315 – 319 nm. Also, the condition of resonance of microcavity is “fair” (“f”) in the wavelength range 320 – 322 nm. The wavelengths of resonance of spherical dielectric microcavity are depicted in Table 8 (Section 6.3.3).

(iii) Spherical dielectric microcavity in the UVA band, with keratinocyte of diameter of 50 µm, has the following parameters:

Core of melanin, radius = 160 nm

Radial dimension of melanin core, r = 160 nm

Refractive index of melanin, n = 1.67

Layer of keratinocyte, radius = 25 µm = 25000 nm

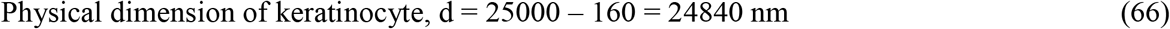

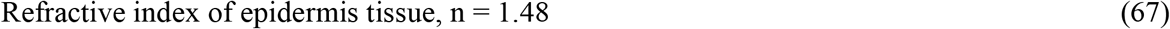

Layer of keratin (physical dimension not needed)

Refractive index of keratin, n = 1.55

The spherical microcavity is said to have been designed for the Bragg wavelength λ_BR_ = 4 x 1.48 x 24840 = 147053 nm (≈ 0.15 mm) in the infrared band of the solar radiation spectrum (from (66) and (67)).

For the keratinocyte of diameter 50 µm, the condition of resonance of microcavity is “good” (“g”) in the wavelength range 328 – 330 nm. Also, the condition of resonance of microcavity is “fair” (“f”) in the wavelength range 331 – 333 nm. The wavelengths of resonance of spherical dielectric microcavity are depicted in Table 8 (Section 6.3.3).

(iv) Spherical dielectric microcavity in the UVA band, with keratinocyte of diameter of 70 µm, has the following parameters:

Core of melanin, radius = 160 nm

Radial dimension of melanin core, r = 160 nm

Refractive index of melanin, n = 1.67

Layer of keratinocyte, radius = 35 µm = 35000 nm

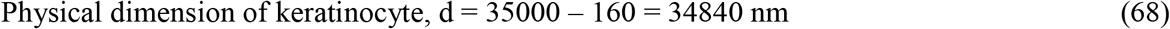

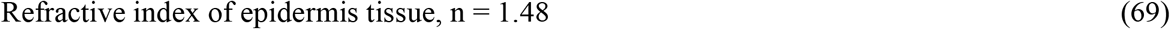

Layer of keratin (physical dimension not needed)

Refractive index of keratin, n = 1.55

The spherical microcavity is said to have been designed for the Bragg wavelength λ_BR_ = 4 x 1.48 x 34840 = 206253 nm (≈ 0.21 mm) in the infrared band of the solar radiation spectrum (from (68) and (69)).

For the keratinocyte of diameter 70 µm, the spherical dielectric microcavity does not reveal any resonance of the spherical dielectric microcavity (“–” in Table 8, Section 6.3.3).

#### 6.3.3 A Summary and Analysis of Wavelengths of Resonance of Spherical Dielectric Microcavity for Brown-Skin Epidermis in the UVB and UVA Bands

The wavelengths of resonance of spherical dielectric microcavity for Brown-skin epidermis in the UVB and UVA bands are given in Table 8.

In the UVB band, in Table 8, the photons of wavelengths 290.0 nm to 297.5 nm are absorbed in the middle and upper portions of the epidermis of Brown-skin. The upper portion of the epidermis absorbs photons of wavelengths 309.5 nm to 314.0 nm. The latter range of propagation wavelengths is due to the underlying approximation of “fair”/ “good” condition of resonance of the spherical dielectric microcavity in the model study. This range of wavelengths is included in the model study. The photons of wavelengths 307.0 nm to 313.0 nm are absorbed in the epidermis close to the stratum corneum. The model propagation ranges of wavelengths are greater than 290 nm for the solar UVB radiation to be effective on the human skin. Also, the model propagation ranges of 290.0 nm to 297.5 nm and 307.0 nm to 314.0 nm are amidst the wavelength range of 290 nm to ∼317 nm of the absorption of UVB photons by 7-dehydrocholesterol in the human skin to synthesize vitamin D_3_ [1]. (The maximal absorption by 7-dehydrocholesterol in the human skin is in the wavelength range 295 nm to 300 nm, peaked around 298 nm.) The model propagation ranges imply the effectiveness of the solar UVB radiation in the wavelength ranges 290 nm to 297.5 nm (≈ 297 nm) and 307 nm to 314 nm for the synthesis of vitamin D_3_ in Brown-skin. The model propagation ranges also imply the application of UVB radiation phototherapies in the wavelength range 290 nm to 297 nm toward synthesis of vitamin D_3_ in Brown-skin (Section 7). The model propagation ranges of 290 nm to 297 nm and 307 nm to 314 nm imply the causative effect of the excessive exposure of solar UVB radiation in the wavelength ranges 290 nm to 297 nm and 307 nm to 314 nm of “delayed” long-term tanning of Brown-skin.

In the UVA band, in Table 8, the photons of wavelengths 315 nm to 342 nm are absorbed in the epidermis of Brown-skin. The model propagation range partly matches the wavelength range of 320 nm to 360 nm that stimulates the production of pigment melanin and skin pigmentation [3]. The model propagation range implies the causative effect of the solar UVA radiation in the wavelength range 315 nm to 342 nm of tanning of Brown-skin, like the skin tanning caused by the UVB radiation (“UVB-like”). The UVA photons of wavelengths 391 nm to 397 nm are absorbed in the epidermis, close to the basal layer, of Brown-skin. The latter narrow range of propagation wavelengths is due to the underlying approximation of “fair”/ “good” condition of resonance of the spherical dielectric microcavity in the model study, and is summarily rejected.

### 6.4 Design and Analysis of a Spherical Dielectric Microcavity in the Epidermis of Black-Skin in the UVB and UVA Bands

A spherical dielectric microcavity, representing an idealized keratinocyte in the epidermis of Black-skin, consists of core of melanin (diameter = 240 nm, radius = 120 nm), and a pair of layers of keratinocyte ((i) diameter = 15 µm, radius = 7.5 µm, (ii) diameter = 30 µm, radius = 15 µm, (iii) diameter = 50 µm, radius = 25 µm, (iv) diameter = 70 µm, radius = 35 µm) and keratin (physical dimension not needed in the computations).

#### 6.4.1 Spherical Dielectric Microcavity for Black-Skin Epidermis in the UVB Band

(i) Spherical dielectric microcavity in the UVB band, with keratinocyte of diameter of 15 µm, has the following parameters (Figure 4):

**Figure 4.**
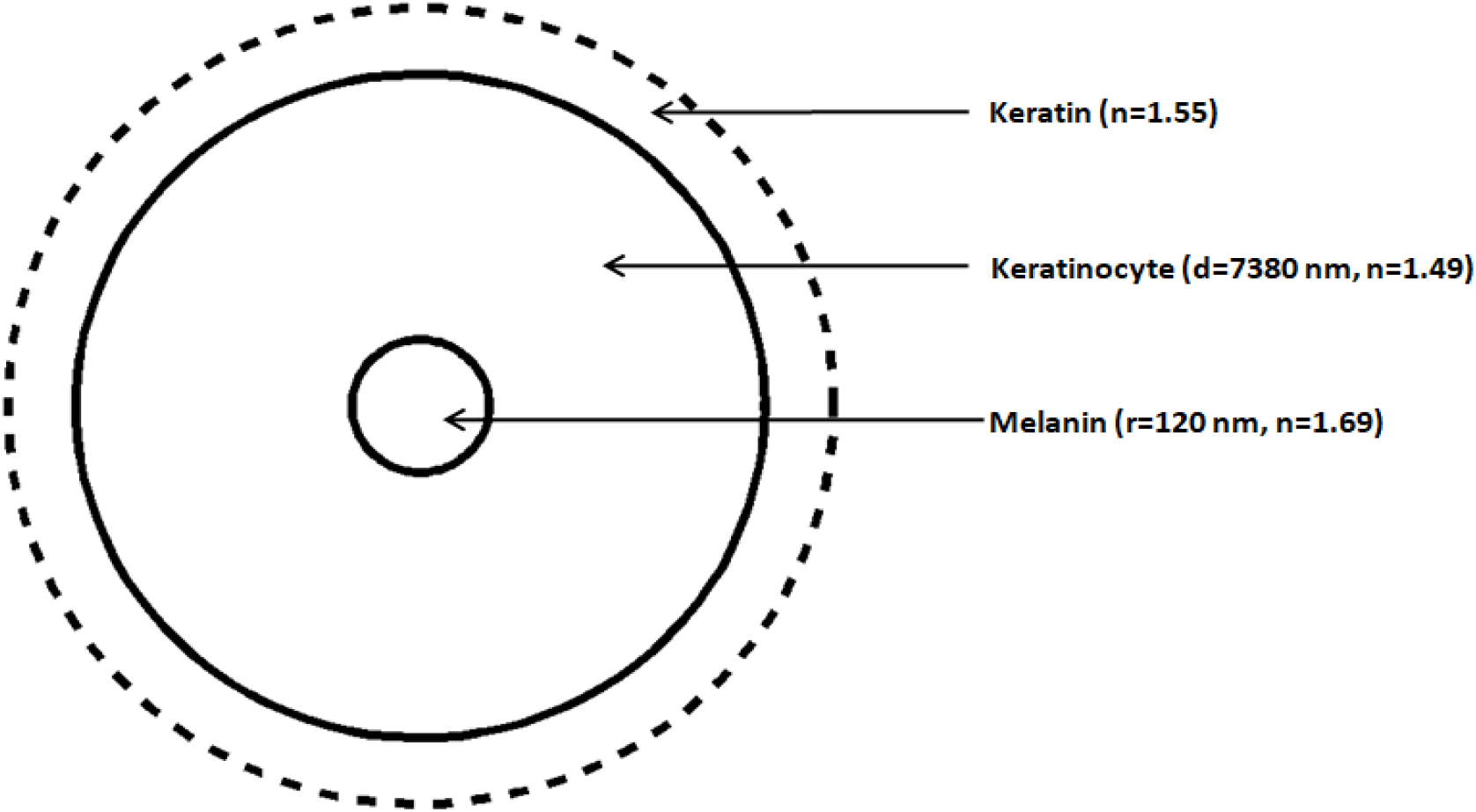
A spherical dielectric microcavity represents a keratinocyte of diameter 15 µm in the epidermis of Black-skin in the UVB band. It consists of core of melanin (radius r = 120 nm, refractive index n = 1.69) and a pair of layers of keratinocyte (physical dimension d = 7380 nm, refractive index n = 1.49) and keratin (dotted, refractive index n = 1.55).

Core of melanin, radius = 120 nm

Radial dimension of melanin core, r = 120 nm

Refractive index of melanin, n = 1.69

Layer of keratinocyte, radius = 7.5 µm = 7500 nm

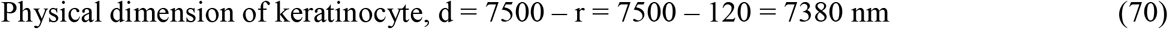

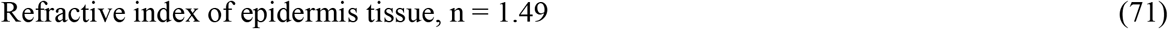

Layer of keratin (physical dimension not needed, dotted in Figure 4)

Refractive index of keratin, n = 1.55

The spherical microcavity is said to have been designed for the Bragg wavelength λ_BR_ = 4 x 1.49 x 7380 = 43985 nm (≈ 0.044 mm) in the infrared band of the solar radiation spectrum (from (70) and (71)).

For the keratinocyte of diameter 15 µm, the spherical dielectric microcavity does not reveal any resonance of the spherical dielectric microcavity (“–” in Table 9, Section 6.4.3).

**Table 9.**
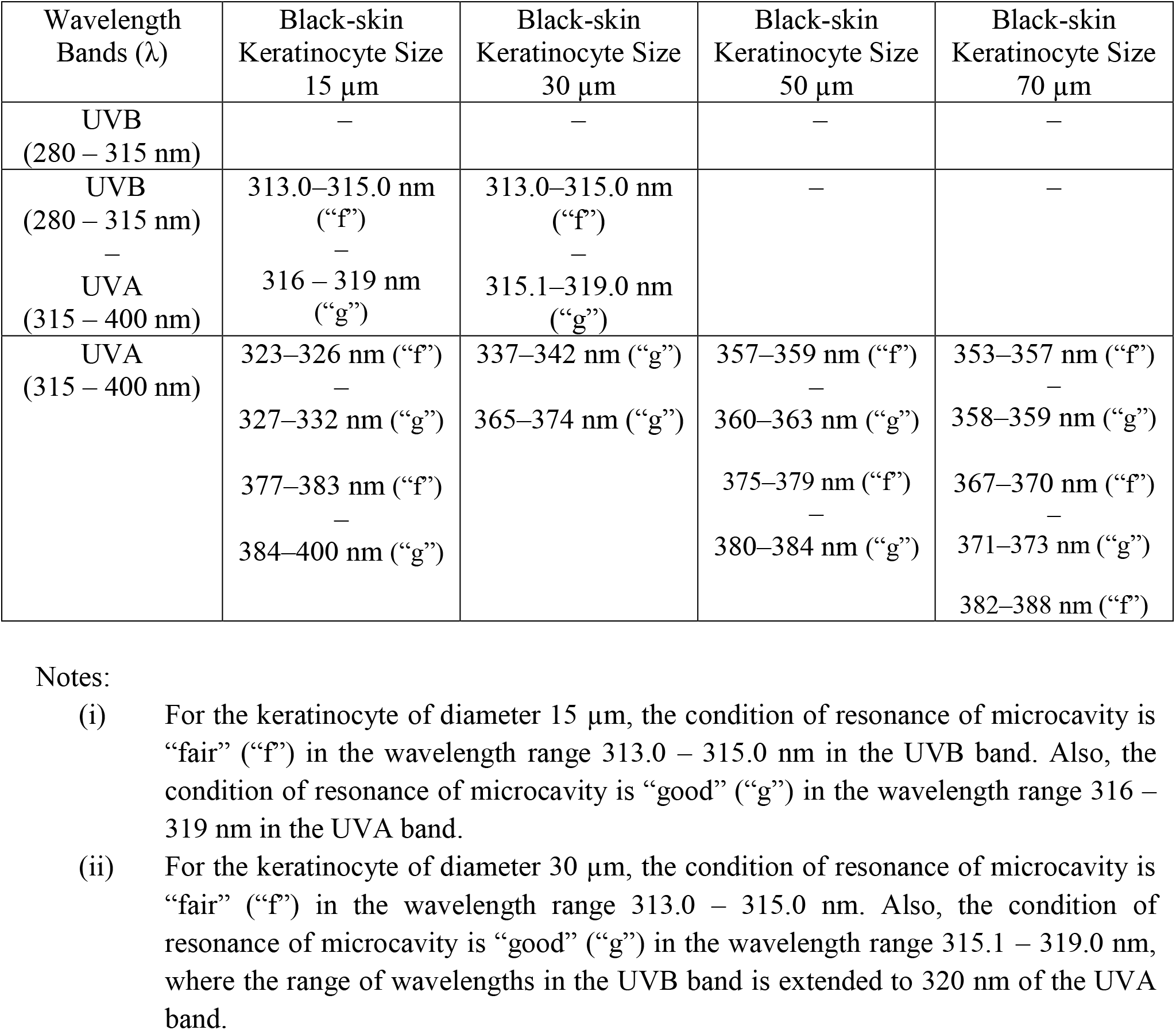
The wavelengths of resonance of spherical dielectric microcavity representing melanin embedded keratinocyte of size 15 µm to 70 µm in the epidermis of Black-skin in the UVB and UVA bands. The condition of resonance is termed as “good” (“g”) or “fair” (“f”). A range of 3 nm in the UVB and 5 nm in the UVA is considered as the wavelengths of resonance. A spherical dielectric microcavity not resonating at a keratinocyte size is represented by “–”

(ii) Spherical dielectric microcavity in the UVB band, with keratinocyte of diameter of 30 µm, has the following parameters:

Core of melanin, radius = 120 nm

Radial dimension of melanin core, r = 120 nm

Refractive index of melanin, n = 1.69

Layer of keratinocyte, radius = 15 µm = 15000 nm

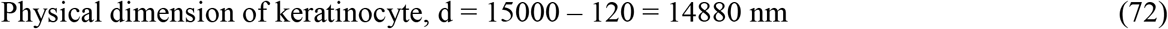

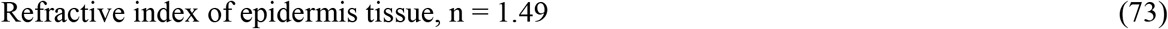

Layer of keratin (physical dimension not needed)

Refractive index of keratin, n = 1.55

The spherical microcavity is said to have been designed for the Bragg wavelength λ_BR_ = 4 x 1.49 x 14880 = 88685 nm (≈ 0.089 mm) in the infrared band of the solar radiation spectrum (from (72) and (73)).

For the keratinocyte of diameter 30 µm, the spherical dielectric microcavity does not reveal any resonance of the spherical dielectric microcavity (“–” in Table 9, Section 6.4.3).

(iii) Spherical dielectric microcavity in the UVB band, with keratinocyte of diameter of 50 µm, has the following parameters:

Core of melanin, radius = 120 nm

Radial dimension of melanin core, r = 120 nm

Refractive index of melanin, n = 1.69

Layer of keratinocyte, radius = 25 µm = 25000 nm

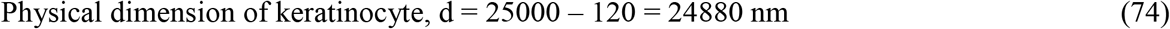

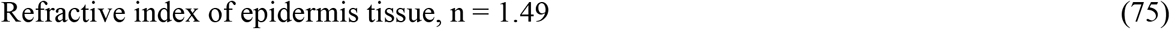

Layer of keratin (physical dimension not needed)

Refractive index of keratin, n = 1.55

The spherical microcavity is said to have been designed for the Bragg wavelength λ_BR_ = 4 x 1.49 x 24880 = 148285 nm (≈ 0.15 mm) in the infrared band of the solar radiation spectrum (from (74) and (75)).

For the keratinocyte of diameter 50 µm, the spherical dielectric microcavity does not reveal any resonance of the spherical dielectric microcavity (“–” in Table 9, Section 6.4.3).

(iv) Spherical dielectric microcavity in the UVB band, with keratinocyte of diameter of 70 µm, has the following parameters:

Core of melanin, radius = 120 nm

Radial dimension of melanin core, r = 120 nm

Refractive index of melanin, n = 1.69

Layer of keratinocyte, radius = 35 µm = 35000 nm

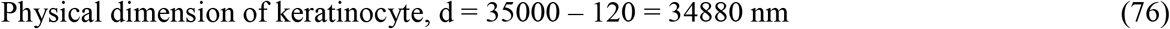

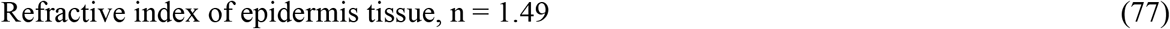

Layer of keratin (physical dimension not needed)

Refractive index of keratin, n = 1.55

The spherical microcavity is said to have been designed for the Bragg wavelength λ_BR_ = 4 x 1.49 x 34880 = 207885 nm (≈ 0.21 mm) in the infrared band of the solar radiation spectrum (from (76) and (77)).

For the keratinocyte of diameter 70 µm, the spherical dielectric microcavity does not reveal any resonance of the spherical dielectric microcavity (“–” in Table 9, Section 6.4.3).

#### 6.4.2 Spherical Dielectric Microcavity for Black-Skin Epidermis in the UVA Band

(i) Spherical dielectric microcavity in the UVA band, with keratinocyte of diameter of 15 µm, has the following parameters:

Core of melanin, radius = 120 nm

Radial dimension of melanin core, r = 120 nm

Refractive index of melanin, n = 1.67

Layer of keratinocyte, radius = 7.5 µm = 7500 nm

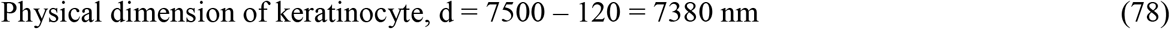

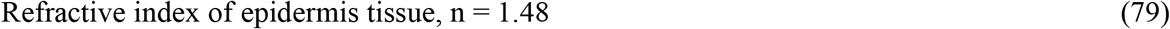

Layer of keratin (physical dimension not needed)

Refractive index of keratin, n = 1.55

The spherical microcavity is said to have been designed for the Bragg wavelength λ_BR_ = 4 x 1.48 x 7380 = 43690 nm (≈ 0.044 mm) in the infrared band of the solar radiation spectrum (from (78) and (79)).

For the keratinocyte of diameter 15 µm: (a) The condition of resonance of microcavity is “fair” (“f”) in the wavelength range 323 – 326 nm. Also, the condition of resonance of microcavity is “good” (“g”) in the wavelength range 327 – 332 nm. (b) The condition of resonance of microcavity is similar to that of “high” (“h”) resonance (Section 6). The condition of resonance of microcavity is “fair” (“f”) in the wavelength range 377 – 383 nm. Also, the condition of resonance of microcavity is “good” (“g”) in the wavelength range 384 – 400 nm. The wavelengths of resonance of spherical dielectric microcavity are depicted in Table 9 (Section 6.4.3).

(ii) Spherical dielectric microcavity in the UVA band, with keratinocyte of diameter of 30 µm, has the following parameters:

Core of melanin, radius = 120 nm

Radial dimension of melanin core, r = 120 nm

Refractive index of melanin, n = 1.67

Layer of keratinocyte, radius = 15 µm = 15000 nm

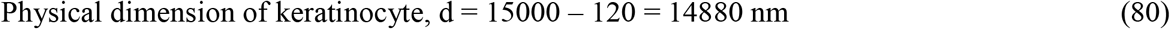

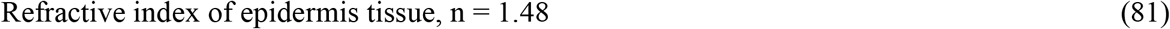

Layer of keratin (physical dimension not needed)

Refractive index of keratin, n = 1.55

The spherical microcavity is said to have been designed for the Bragg wavelength λ_BR_ = 4 x 1.48 x 14880 = 88090 nm (≈ 0.088 mm) in the infrared band of the solar radiation spectrum (from (80) and (81)).

For the keratinocyte of diameter 30 µm: (a) The condition of resonance of microcavity is “good” (“g”) in the wavelength range 337 – 342 nm. (b) The condition of resonance of microcavity is similar to that of “high” (“h”) resonance (Section 6). The condition of resonance of microcavity is “good” (“g”) in the wavelength range 365 – 374 nm. The wavelengths of resonance of spherical dielectric microcavity are depicted in Table 9 (Section 6.4.3).

(iii) Spherical dielectric microcavity in the UVA band, with keratinocyte of diameter of 50 µm, has the following parameters:

Core of melanin, radius = 120 nm

Radial dimension of melanin core, r = 120 nm

Refractive index of melanin, n = 1.67

Layer of keratinocyte, radius = 15 µm = 25000 nm

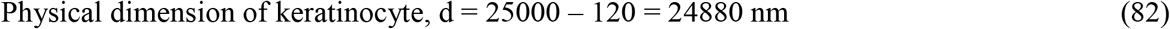

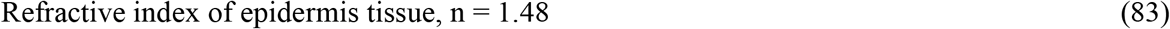

Layer of keratin (physical dimension not needed)

Refractive index of keratin, n = 1.55

The spherical microcavity is said to have been designed for the Bragg wavelength λ_BR_ = 4 x 1.48 x 24880 = 147290 nm (≈ 0.15 mm) in the infrared band of the solar radiation spectrum (from (82) and (83)).

For the keratinocyte of diameter 50 µm: (a) The condition of resonance of microcavity is similar to that of “high” (“h”) resonance (Section 6). The condition of resonance of microcavity is “fair” (“f”) in the wavelength range 357 – 359 nm. Also, the condition of resonance of microcavity is “good” (“g”) in the wavelength range 360 – 363 nm. (b) The condition of resonance of microcavity is similar to that of “high” (“h”) resonance (Section 6). The condition of resonance of microcavity is “fair” (“f”) in the wavelength range 375 – 379 nm. Also, the condition of resonance of microcavity is “good” (“g”) in the wavelength range 380 – 384 nm. The wavelengths of resonance of spherical dielectric microcavity are depicted in Table 9 (Section 6.4.3).

(iv) Spherical dielectric microcavity in the UVA band, with keratinocyte of diameter of 70 µm, has the following parameters:

Core of melanin, radius = 120 nm

Radial dimension of melanin core, r = 120 nm

Refractive index of melanin, n = 1.67

Layer of keratinocyte, radius = 15 µm = 35000 nm

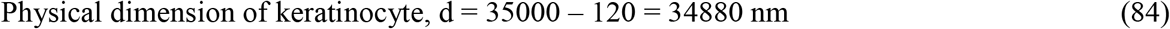

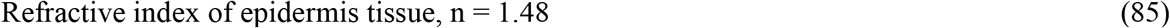

Layer of keratin (physical dimension not needed)

Refractive index of keratin, n = 1.55

The spherical microcavity is said to have been designed for the Bragg wavelength λ_BR_ = 4 x 1.48 x 34880 = 206490 nm (≈ 0.15 mm) in the infrared band of the solar radiation spectrum (from (84) and (85)).

For the keratinocyte of diameter 70 µm: (a) The condition of resonance of microcavity is similar to that of “high” (“h”) resonance (Section 6). The condition of resonance of microcavity is “fair” (“f”) in the wavelength range 353 – 357 nm. Also, the condition of resonance of microcavity is “good” (“g”) in the wavelength range 358 – 359 nm. (b) The condition of resonance of microcavity is similar to that of “high” (“h”) resonance (Section 6). The condition of resonance of microcavity is “fair” (“f”) in the wavelength range 367 – 370 nm. Also, the condition of resonance of microcavity is “good” (“g”) in the wavelength range 371 – 373 nm. The wavelengths of resonance of microcavity are depicted in Table 9. (c) The condition of resonance of microcavity is similar to that of “high” (“h”) resonance (Section 6). The condition of resonance of microcavity is “fair” (“f”) in the wavelength range 382 – 388 nm. The wavelengths of resonance of spherical dielectric microcavity are depicted in Table 9 (Section 6.4.3).

#### 6.4.3 A Summary and Analysis of Wavelengths of Resonance of Spherical Dielectric Microcavity for Black-Skin Epidermis in the UVB and UVA Bands

The wavelengths of resonance of spherical dielectric microcavity for Black-skin epidermis in the UVB and UVA bands are given in Table 9.

In the hybrid (UVB – UVA) band, in Table 9, the photons of wavelengths 313 nm to 319 nm are absorbed in the epidermis, close to the basal layer and the middle portion, of Black-skin. The model propagation range of wavelengths is greater than 290 nm for the solar (UVB – UVA) radiation to be effective on the human skin. Also, a part of the model propagation range of 313 nm to ∼317 nm is at the rear end of the wavelength range of 290 nm to ∼317 nm of the absorption of (UVB – UVA) photons by 7-dehydrocholesterol in the human skin to synthesize vitamin D_3_ [1]. The model propagation range implies the effectiveness of the solar (UVB – UVA) radiation in the wavelength range 313 nm to ∼317 nm for the synthesis of vitamin D_3_ in Black-skin. The model propagation range also implies the application of (UVB – UVA) radiation phototherapies in the wavelength range 313 nm to ∼317 nm toward synthesis of vitamin D_3_ in Black-skin (Section 7). The model propagation range of 313 nm to 319 nm implies the causative effect of the solar (UVB – UVA) radiation in the wavelength range 313 nm to 319 nm of tanning of Black-skin.

In the UVA band, in Table 9, the photons of wavelengths 353 nm to 400 nm are absorbed in the epidermis of Black-skin. The model propagation range encompasses the peak range of tanning of skin of 340 nm to 370 nm [7, Review]. The model propagation range implies the causative effect of the solar UVA radiation in the wavelength range 353 nm to 400 nm of tanning of Black-skin. The UVA photons of wavelengths 323 nm to 332 nm and 337 nm to 342 nm are absorbed in the epidermis, close to the basal layer and the middle portion, respectively, of Black-skin. The latter narrow ranges of propagation wavelengths is due to the underlying approximation of “fair”/ “good” condition of resonance of the spherical dielectric microcavity in the model study, and are summarily rejected.

## 7. Application of Model Wavelengths of Propagation of Electromagnetic Waves in the Epidermis of White-Skin, Pale Yellow-Skin, Brown-Skin and Black-Skin

The propagation of electromagnetic waves in the UVB (280 – 315 nm) and UVA (315 – 400 nm) bands in the epidermis of a skin type finds the following applications: (a) Synthesis of vitamin D_3_ – solar radiation induced, (b) Synthesis of vitamin D_3_ – phototherapy, (c) Skin tanning – solar and phototherapy (therapeutic use in skin care), and (d) Preparation of sunscreen gel. The phototherapy for the synthesis of vitamin D_3_ is necessities to individuals who are allergic to the medicinal form of vitamin D_3_, and are solely dependent the on solar UVB radiation. The model (model computed) wavelengths of these applications are summarized in Table 10, which are based on the wavelengths of absorption of photons in the epidermis of the four skin types in Table 6 – Table 9. The model computations of the wavelengths of propagation in the White-skin set the efficacy of the model study.

**Table 10.**
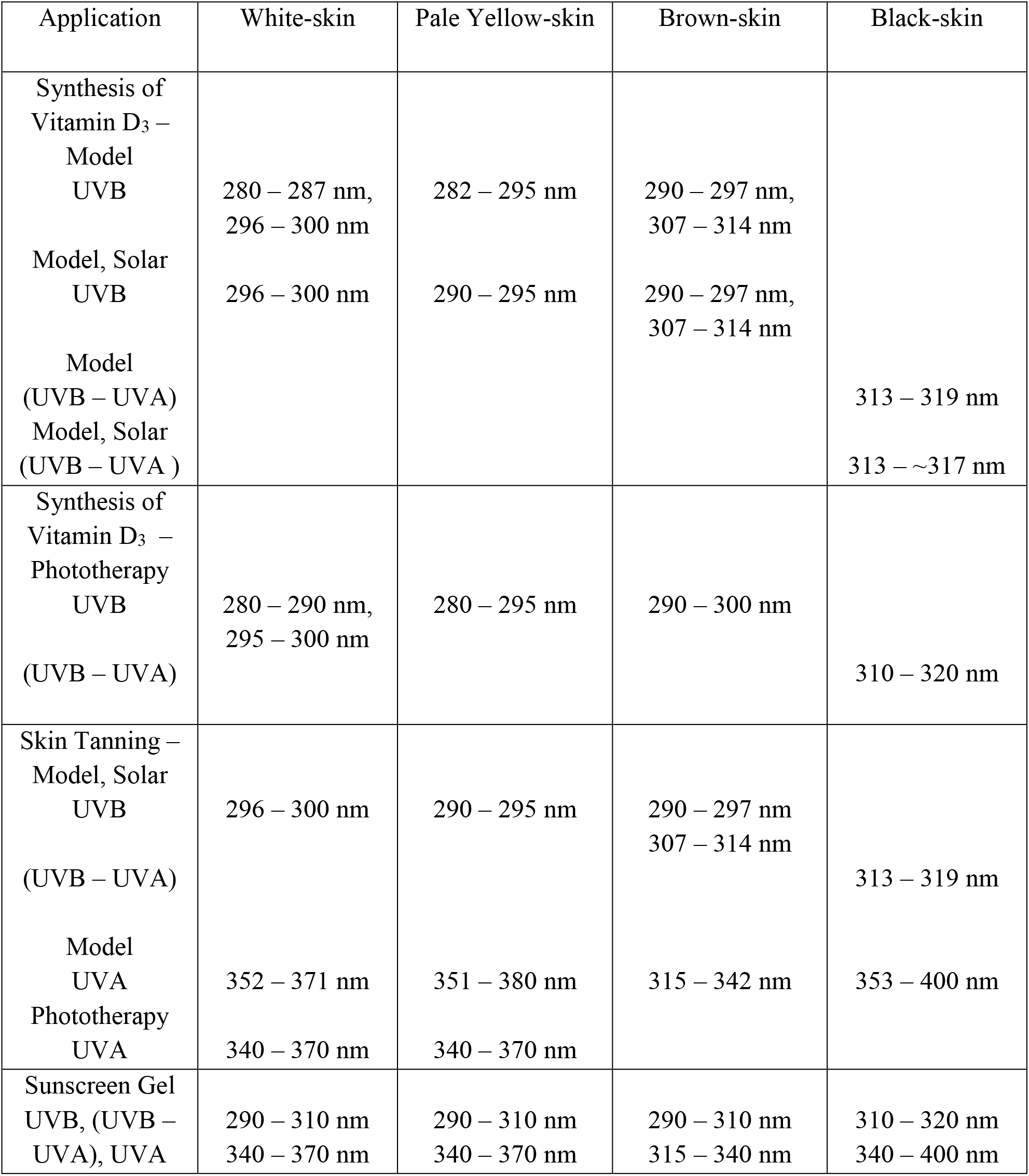
Summary of wavelengths of propagation in the UVB (280 – 315 nm) and UVA (315 – 400 nm) bands in the epidermis of White-skin, Pale Yellow-skin, Brown-skin and Black-skin

The wavelength ranges of the aforementioned applications in the UVB (280 – 315 nm) and UVA (315 – 400 nm) bands for the four skin types are described below. The following statements are applicable to various skin types. The solar UVB radiation longer than 290 nm alone reaches the earth’s surface, since the UVB wavelengths shorter than 290 nm are absorbed by the atmosphere. Prolonged sun exposure is required in pigmented skin to effectively synthesize vitamin D_3_ [25]. The tanning caused by the solar UVA radiation does not protect the skin from the sunburn (erythema) by caused by the UVB radiation [7, Review]. The photochemical mechanism of tanning caused by the solar UVA radiation in the wavelength range 320 nm to 360 nm is similar to that of the solar UVB radiation (“UVB-like”), though less effectively [3]. The tanning caused by the solar UVB/UVA(320 – 360 nm) radiation protects the skin from the sunburn (erythema). The sunscreen gel drastically diminishes the effectiveness of the solar UVB radiation in order to synthesize vitamin D_3_ in the skin.

(i) White-skin
  a. The solar UVB radiation in the model wavelength range 296 nm to 300 nm is effective for the synthesis of vitamin D_3_. The wavelength range for the maximal synthesis of vitamin D_3_ by the compound 7-dehydrocholesterol in skin cells is 295 nm to 300 nm, with peak synthesis around 298 nm [1]. The effective wavelength range for the solar induced synthesis of vitamin D_3_ is 290 nm to 310 nm [1].
  b. The UVB radiation in the model wavelength range 280 nm to 287 nm (≈ 290 nm) may be applied toward phototherapies for the synthesis of vitamin D_3_. The wavelength range is apart from the wavelength range 296 nm (≈ 295 nm) to 300 nm for the synthesis of vitamin D_3_. The human skin is prone to sunburn (erythema) at wavelengths less than 290 nm over excessive UVB exposure [7, Review].
  c. The solar UVB radiation in the wavelength range 296 nm to 300 nm causes “delayed” long-term tanning of skin over excessive exposure. The solar UVA radiation in the model wavelength range 352 nm to 371 nm causes “immediate” tanning of skin. The UVA range of wavelengths 352 (≈ 340) nm to 371 (≈ 370) nm may be applied toward phototherapies for “immediate” tanning of skin. The wavelength range for the peak tanning of skin is around 340 nm to 370 nm [7, Review].
  d. A sunscreen gel may contain organic (aromatic) compounds to absorb the solar UVB radiation of wavelengths 290 nm to 310 nm and the solar UVA radiation of wavelengths 340 nm to 370 nm, and convert the UV (UVB, UVA) radiation to the IR radiation (heat).
(ii) Pale Yellow-skin
  a. The solar UVB radiation in the model wavelength range 290 nm to 295 nm is effective for the synthesis of vitamin D_3_, where the peak synthesis of vitamin D_3_ is around 298 nm [1]. The effective synthesis of vitamin D_3_ is in the wavelength range 290 nm to 310 nm [1]. The effectiveness of synthesis of vitamin D_3_ in Pale Yellow-skin is to a lesser extent compared to White-skin.
  b. The UVB radiation in the model wavelength range 282 nm (≈ 280 nm) to 295 nm may be applied toward phototherapies for the synthesis of vitamin D_3_. The human skin may be subject to sunburn (erythema) at wavelengths less than 290 nm [7, Review].
  c. The solar UVB radiation in the wavelength range 290 nm to 295 nm causes “delayed” long-term tanning of skin over excessive exposure. The solar UVA radiation in the model wavelength range 351 nm to 380 nm causes “immediate” tanning of skin. The UVA range of wavelengths 351 nm (≈ 340 nm) to 380 nm (≈ 370 nm) may be applied toward phototherapies for “immediate” tanning of skin. The peak tanning of skin is around wavelengths 340 nm to 370 nm [7, Review].
  d. A sunscreen gel for the Pale Yellow-skin may contain organic compounds to absorb the solar UVB and UVA radiation of wavelengths 290 nm to 310 nm and 340 to 370 nm, respectively, as in the case of White-skin.
(iii) Brown-skin
  a. The solar UVB radiation in the model wavelength ranges 290 nm to 297 nm and 307 nm to 314 nm are effective for the synthesis of vitamin D_3_, where the peak synthesis of vitamin D_3_ is around 298 nm [1]. The effective synthesis of vitamin D_3_ is in the wavelength range 290 nm to 310 nm, where the overall synthesis of vitamin D_3_ in the skin is in the wavelengths 290 – ∼317 nm [1]. Apparently, the solar UVB radiation is more effective to synthesize vitamin D_3_ in Brown-skin compared to Pale Yellow-skin. The Brown-skin being prone to pigmentation over exposure to sun, the solar UVB radiation is less effective to synthesize vitamin D_3_ in Brown-skin compared to Pale Yellow-skin.
  b. The UVB radiation in the model wavelength range 290 nm to 297 nm (≈ 300 nm) may be applied toward phototherapies for the synthesis of vitamin D_3_.
  c. The solar UVB radiation in the wavelength ranges 290 nm to 297 nm (≈ 300 nm) and 307 nm (≈ 310 nm) to 314 nm (≈ 315 nm) causes “delayed” long-term tanning of skin over excessive exposure. The solar UVA radiation in the model wavelength range 315 nm to 342 nm (≈ 340 nm) causes tanning of skin, where the wavelength range of “UVB-like” tanning of skin is around 320 nm to 360 nm [3]. Consequently, the Brown-skin is prone to pigmentation and is less effective to synthesize vitamin D_3_.
  d. A sunscreen gel for the Brown-skin may contain organic compounds to absorb the solar UVB and UVA radiation of wavelengths 290 nm to 310 nm and 315 nm to 340 nm, respectively.
(iv) Black-skin
  a. The solar (UVB – UVA) radiation in the wavelength range 313 nm to ∼317 nm is effective for the synthesis of vitamin D_3._ Overall, the epidermis layer of the skin synthesizes vitamin D_3_ in the wavelengths 290 – ∼317 nm [1]. The Black-skin being least effective, prolonged sun exposure is required to effectively synthesize vitamin D_3_.
  b. The (UVB – UVA) radiation in the wavelength range 313 nm (≈ 310 nm) to ∼317 nm (≈ 320 nm) may be applied toward phototherapies for the synthesis of vitamin D_3_. Prolonged phototherapy is required to effectively synthesize vitamin D_3_.
  c. The solar (UVB – UVA) radiation in the wavelength range 313 nm (≈ 310 nm) to 319 nm (≈ 320 nm) causes tanning of skin. The solar UVA radiation in the model wavelength range 353 nm (≈ 340 nm) to 400 nm also causes tanning of skin, where the peak tanning of skin is around 340 nm to 370 nm [7, Review].
  d. A sunscreen gel for the Black-skin may contain organic compounds to absorb the solar (UVB – UVA) and UVA radiation of wavelengths 310 nm to 320 nm and 340 nm to 400 nm, respectively.

Some of the well-known facts to have sufficient levels of vitamin D_3_, as a rule of thumb, not discussed in the study include exposure of about 15% to 20% of body to sun for about 15 minutes to 20 minutes in the open sunshine. Moreover, the exposure may be for about 2 hours before local noon in summer and for about 2 hours around local noon in winter. These are the time periods when the sun is close to being overhead and the sun’s rays of intense solar UVB radiation fall almost straight on earth. Also, the atmospheric ozone absorbs the UVB radiation to the minimum. There are environmental parameters, like, (glass) windowpane, pollution, etcetera, that block the UVB radiation that synthesizes vitamin D_3_ in skin.

## 8. Conclusion

The study presents model computations on the role of ultraviolet (UV) radiation toward (i) synthesis of vitamin D_3_ in skin, (ii) therapeutic use in skin tanning and (iii) preparation of sunscreen gel for skin protection over sun exposure. The study computationally evaluates the documented results that the solar UVB radiation, to synthesize vitamin D_3_, is the most effective in White-skin and Pale Yellow-skin, and the least effective in Black-skin. The solar UVB radiation also synthesizes sufficient levels of vitamin D_3_ in Brown-skin. It is established that a single sunscreen gel for skin protection may not suffice the very purpose for different skin types.

Model computations of the wavelengths of propagation of electromagnetic waves in the ultraviolet (UV) band in the epidermis cells keratinocytes of skin are performed. The four skin types are: (a) White-skin, (b) Pale Yellow-skin, (c) Brown-skin and (d) Black-skin. An ideal spherical keratinocyte in the epidermis is considered, with a spherical melanin granule at the center and a layer of keratin molecules surrounding the cell. Such an idealized keratinocyte represents a spherical dielectric microcavity, with the published data on refractive indices of epidermis tissue, melanin and keratin. The thickness of layer of keratin does not occur in the computations. The sizes of keratinocyte cells and skin type specific-melanin granules are estimated in the present study. The model computations are carried out in four keratinocyte sizes, 15 µm to 70 µm. The calculated sizes of melanin granules are: (a) White-skin - 800 nm, (b) Pale Yellow-skin - 460 nm, (c) Brown-skin - 320 nm and (d) Black-skin - 240 nm. Electromagnetic waves are incident in the core melanin of the spherical microcavity. The range of wavelengths of resonance of the microcavity is the model (model computed) wavelength range of propagation, where photons are absorbed in the cells keratinocytes.

In (a) White-skin, the model UVB radiation wavelength ranges are approximately 280 – 290 nm and 295 – 300 nm. The UVB radiation wavelength range 295 – 300 nm is observed in the epidermis cells keratinocytes of size 15 µm, which are in the vicinity of the basal layer. The solar UVB radiation in the model wavelength range maximally synthesizes vitamin D_3_ in White-skin. The UVB radiation in the wavelength range 280 – 290 nm and 295 – 300 nm may be applied toward phototherapies for the synthesis of vitamin D_3_. However, the human skin is prone to sunburn (erythema) at wavelengths less than 290 nm. The solar UVB radiation in the model wavelength range leads to the solar UVB-caused tanning. The model UVA radiation wavelength range of approximately 340 – 370 nm leads to the solar UVA-caused tanning of skin; which is the documented wavelength range for the peak tanning of skin. Thus, the UVA radiation phototherapies toward tanning of skin may be applied in the wavelength range of 340 – 370 nm. A sunscreen gel may contain organic compounds to absorb the effective solar UVB and the UVA radiation of wavelengths of 290 – 310 nm and 340 – 370 nm, respectively, in White-skin.

In (b) Pale Yellow-skin, the model UVB radiation wavelength range is approximately 280 – 295 nm. The UVB radiation wavelength range 290 – 295 nm is observed in the epidermis cells keratinocytes of size 30 µm and 50 µm, which are farther from the basal layer toward the stratum corneum. The solar UVB radiation in the wavelength range of 290 – 295 nm synthesizes vitamin D_3_ to high levels in Pale Yellow-skin. The UVB radiation in the wavelength range 280 – 295 nm may be applied toward phototherapies for the synthesis of vitamin D_3_. The UVB radiation wavelength range of 290 – 295 nm leads to the solar UVB-caused tanning of skin. The model UVA radiation wavelength range of approximately 340 – 370 nm leads to the solar UVA-caused tanning of skin. The UVA radiation phototherapies toward tanning of skin may be applied in the wavelength range of 340 – 370 nm, as in the case of White-skin. A sunscreen gel may be prepared to absorb the effective solar UVB and the UVA radiation wavelengths of 290 – 310 nm and 340 – 370 nm, respectively, as in the case of White-skin.

In (c) Brown-skin, the model UVB radiation wavelength ranges are 290 – 297 nm and 307 – 314 nm. The dominant UVB radiation wavelength range 290 – 297 nm is observed in the epidermis cells keratinocytes of size 30 µm and 50 µm, which are farther from the basal layer toward the stratum corneum, as in the case of Pale Yellow-skin. The solar UVB radiation in the two ranges of wavelength together synthesizes vitamin D_3_ in Brown-skin. The Brown-skin being prone to pigmentation, the solar UVB radiation induces ample levels of vitamin D_3_ in Brown-skin. The UVB radiation in the wavelength range 290 – 300 nm may be applied toward phototherapies for the synthesis of vitamin D_3_. The model UVB radiation wavelength ranges in the two wavelength ranges together leads to the solar UVB-caused tanning of skin. The model UVA radiation wavelength range of approximately 315 – 340 nm leads to the solar UVA-caused tanning of skin, which is “UVB-like” tanning of skin. The tannings caused by the UVB and UVA radiation make the Brown-skin prone to pigmentation over exposure to sun. A sunscreen gel may be prepared to absorb the effective solar UVB and the UVA radiation wavelengths of 290 – 310 nm and 315 – 340 nm, respectively, in Brown-skin.

In (d) Black-skin, the model (UVB – UVA) radiation wavelength range is 313 – 319 nm. The (UVB – UVA) radiation wavelength range 313 – ∼317 nm is observed in the epidermis cells keratinocytes of size 15 µm and 30 µm, which are ‘together’ nearer the basal layer, as in the case of White-skin. The solar (UVB – UVA) radiation in the wavelength range 313 – ∼317 nm is the least effective to synthesize vitamin D_3_ in Black-skin. The (UVB – UVA) radiation in the wavelength range 310 – 320 nm may be applied toward phototherapies for the synthesis of vitamin D_3_. Prolonged phototherapy is required to effectively synthesize vitamin D_3_. The model (UVB – UVA) radiation wavelength range leads to the solar (UVB – UVA)-caused tanning of skin. The model UVA radiation wavelength range of approximately 340 – 400 nm leads to the solar UVA-caused tanning of skin. A sunscreen gel may be prepared to absorb the effective solar (UVB – UVA) and the UVA radiation wavelengths of 310 – 320 nm and 340 – 400 nm, respectively, in Black-skin.

The future studies are focused on the model computations of propagation of electromagnetic waves in the visible (400 – 700 nm) and part of near-infrared (700 – 800 nm) bands in the epidermis of the four skin types, considered in the present study.

## Appendix A

The wavelength (λ, nm) dependence of the real refractive index (n_r_) of epidermis tissue is fitted with the Cauchy dispersion relation,

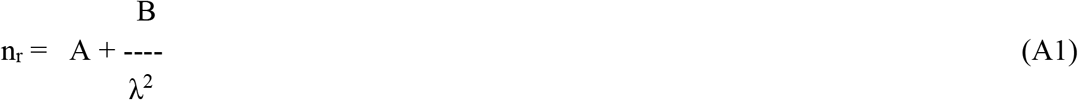

where the fitted parameters A and B are obtained by the method of non-linear least-squares [26]. A smoothened curve is traced between n_r_ versus λ (nm) given in Figure 6 of Ding et al. [19]. Eleven values of n_r_ to 3-decimal points are obtained between 325 nm and 800 nm. (The data need not be equally spaced for the purpose of least-squares fitting.) A computer program in C is written that uses the ‘function’ FCHISQ (page 194) and ‘subroutine’ GRIDLS (page 213) in FORTRAN of Bevington [26]. (The computer program is also suitable for fitting linear functions to data.) The optimum values of the parameters A and B so-obtained are 1.415 and 8.018 × 10^3^. (In Reference 19, Ding et al. have fitted the Cauchy dispersion relation n_r_ = A + B/λ^2^ + C/λ^4^ to the data between n_r_ versus λ for the dermis tissue. The fitted values of the parameters A, B and C are 1.3696, 3.9168 × 10^3^ and 2.5588 × 10^3^, respectively, for the dermis tissue [19, Table 2].)

The wavelength (λ, nm) dependence of the refractive index (n) of melanin is fitted with the Cauchy dispersion relation (A1) by the method of non-linear least-squares. A smoothened curve is traced between n versus λ (nm) in Figure 2 of Repenko et al. [20]. Twelve values of n to 2-decimal points are obtained between 250 nm and 800 nm. The computer program in C for the non-linear least-squares fitting gives the optimum values of the parameters A and B as 1.605 and 9.251 × 10^3^, respectively.

## Acknowledgements

The authors sincerely thank Dr. P. N. Vijaykumar, formerly of the CSIR-National Physical Laboratory, for his valuable comments and insights during the course of the study. The authors express gratitude to CSIR for providing the necessary facilities to carry out this work.

